# Hypoxia-inducing cryogels uncover key cancer-immune cell interactions in an oxygen-deficient tumor microenvironment

**DOI:** 10.1101/2023.01.10.523477

**Authors:** Thibault Colombani, Zachary J. Rogers, Khushbu Bhatt, James Sinoimeri, Lauren Gerbereux, Mohammad Hamrangsekachaee, Sidi A. Bencherif

**Affiliations:** Department of Chemical Engineering, Northeastern University, Boston, MA 02115, United States of America; Department of Pharmaceutical Sciences, Northeastern University, Boston, MA, United States; Department of Bioengineering, Northeastern University, Boston, MA 02115, United States of America; Harvard John A. Paulson School of Engineering and Applied Sciences, Harvard University, Cambridge, MA 02138, United States of America; Biomechanics and Bioengineering (BMBI), UTC CNRS UMR 7338, University of Technology of Compiègne, Sorbonne University, 60203 Compiègne, France

**Keywords:** cryogels, hypoxia, cancer, immunosuppression, drug resistance

## Abstract

Hypoxia, an important feature of solid tumors, is a major factor shaping the immune landscape, and several cancer models have been developed to emulate hypoxic tumors. However, to date, they still have several limitations, such as the lack of reproducibility, inadequate biophysical cues, limited immune cell infiltration, and poor oxygen (O_2_) control, leading to non-pathophysiological tumor responses. As a result, it is essential to develop new and improved cancer models that mimic key features of the tumor extracellular matrix and recreate tumor-associated hypoxia while allowing cell infiltration and cancer-immune cell interactions. Herein, hypoxia-inducing cryogels (HICs) have been engineered using hyaluronic acid (HA) as macroporous scaffolds to fabricate three-dimensional microtissues and model a hypoxic tumor microenvironment. Specifically, tumor cell-laden HICs have been designed to deplete O_2_ locally and induce long-standing hypoxia. This state of low oxygen tension, leading to HIF-1α stabilization in tumor cells, resulted in changes in hypoxia-responsive gene expression and phenotype, a metabolic adaptation to anaerobic glycolysis, and chemotherapy resistance. Additionally, HIC-supported tumor models induced dendritic cell (DC) inhibition, revealing a phenotypic change in plasmacytoid B220^+^ DC (pDC) subset and an impaired conventional B220^−^ DC (cDC) response in hypoxia. Lastly, our HIC-based melanoma model induced CD8+ T cell inhibition, a condition associated with the downregulation of pro-inflammatory cytokine secretion, increased expression of immunomodulatory factors, and decreased degranulation and cytotoxic capacity of T cells. Overall, these data suggest that HICs can be used as a tool to model solid-like tumor microenvironments and identify a phenotypic transition from cDC to pDC in hypoxia and the key contribution of HA in retaining cDC phenotype and inducing their hypoxia-mediated immunosuppression. This technology has great potential to deepen our understanding of the complex relationships between cancer and immune cells in low O_2_ conditions and may pave the way for developing more effective therapies.

## Introduction

The tumor microenvironment (TME) plays a decisive role in cancer initiation and dissemination.(*1*) However, despite massive efforts in cancer research, the TME continues to be poorly emulated and understood. Complex by its composition, the TME is often characterized by low O_2_ levels known as hypoxia (∼0.5–3% O_2_),(*2*) a condition modulating several essential components such as the extracellular matrix, bioactive substances (e.g., glycosaminoglycans), tumor cells, and immune cells.(*3*) Hypoxia, a hallmark of aggressive cancers,(*4*) leads to cell metabolic changes and adaptation, tumor cell growth and invasion, resistance to apoptosis, and drug resistance.(*5*) Moreover, the TME has been shown to induce hypoxia-mediated immunosuppression linked to immune evasion.(*6, 7*) Therefore, unraveling the complex relationships between cancer cells and immune cells in hypoxia is critical, not only to enhance chemotherapy,(*8*) radiotherapy,(*9*) and immunotherapy,(*10*) but also to find more effective strategies that can unlock the full potential of the patient’s immune system to fight back against cancer.

Animal models such as rodents have been perceived as an invaluable tool to better understand the role of the hypoxic TME on cancer development and resistance to cancer therapies. However, they are costly, time-consuming, and a poor predictor in human clinical trials.(*11, 12*) Therefore, in recent years, several approaches to modeling solid hypoxic tumors have been developed, ranging from two-dimensional (2D) to three-dimensional (3D) cell culture models.(*13*) Standard cancer models in 2D are convenient and can reproduce hypoxia-driven biochemical cues, but their lack of 3D cellular organization and an extracellular matrix (ECM) result in non-pathophysiological tumor cell responses.(*14*) To overcome these challenges, cell culture models in 3D such as spheroids and organoids have been developed to emulate the complexity of tumor-like architecture and provide biophysical cues.(*15*) Yet, they still fail to recreate the cellular and matrix complexity of native tumor tissues, be reproducible consistently, provide key biomechanical cues, and induce a tumor-like O_2_ deficient environment.(*16, 17*) Therefore, there is a dire need to develop new strategies and create better cancer models. To advance our understanding of cancer immunology, new approaches should be explored to induce consistent and homogeneous tumor hypoxia, provide physical support to cells, offer a porous network in 3D to facilitate cell infiltration and interactions, display key ECM components, and ultimately emulate aspects of the immunosuppressive TME.

Engineering innovative biomaterials to mimic the TME has gained great interest in the field, especially for drug screening, and to better understand cancer biology and immunology.(*18*) For instance, Matrigel and other biomimetic hydrogels composed of collagen and HA have been extensively investigated to recreate the tumor ECM.(*19*) However, such biomaterials do not display several key features of solid tumors, including control over low O_2_ tension.(*20*) Recent studies have shown that O_2_-consuming enzymes can be used as a reliable alternative to induce hypoxia in aqueous solutions, hydrogels, and microfluidic devices.(*21–23*) The most common strategy relies on glucose oxidase (GOX), a protein catalyzing the oxidation of β-D-glucose to D-glucono-δ-lactone from the medium while quickly depleting O_2_.(*24, 25*) Despite efficient induction of hypoxic conditions, these approaches induce short-term hypoxia and can be highly toxic to cells. In addition, standard hydrogels also create a physical barrier due to their mesoporous structure, restricting tumor cell organization and immune cell infiltration as well as the diffusion of solutes (biomolecules, drugs, nutrients, etc.).(*26*) Therefore, designing scaffolds mimicking tumor ECM with a highly interconnected macroporous network is critical to facilitate the diffusion of solutes, allow immune cell infiltration, and promote cell-cell interactions and organization.(*27, 28*) Our team has recently reported the development of innovative macroporous and injectable cryogels as promising biomaterial for tissue engineering, tumor modeling, and immunotherapies.(*29–32*) Cryogels possess unique features such as shame memory properties, high elasticity, and exhibit an interconnected and large porous structure. Furthermore, cryogels have been successfully used as biomimetic scaffolds to engineer tumor microtissues(*33*) and allow easy functionalization with proteins, enzymes, and peptides,(*34*) making them an ideal platform for cancer immunology research.

In this work, we hypothesized that advanced biomaterials can be designed as a platform to develop improved cancer models, recapitulating key aspects of the environmental cues in solid tumors, such as the physicochemical properties of the ECM and a state of oxygen deficiency. Additionally, we postulated that these biomaterials could be used as a tool to better understand cancer immunology under hypoxic conditions. To this end, we fabricated 3D HICs to quickly create an O_2_-deprived environment and induce cellular hypoxia. HICs were constructed with HA, a major component of the tumor ECM that accumulates in the TME in response to hypoxic conditions.(*35*) The cryogels were covalently grafted with GOX and catalase (CAT) to deplete O_2_ rapidly and safely. First, we characterized the physical properties of HICs and then tested their capacity to induce hypoxia (∼1% O_2_) in a controlled and sustained fashion. Next, we investigated the potential of HICs when infused with cancer cells to emulate hypoxia-driven phenotypic changes, metabolic adaptation to anaerobic glycolysis, and chemoresistance. Finally, we evaluated the effect of HICs on immune cells such as dendritic cells (DCs), principal antigen-presenting cells and major regulator of the adaptive immune system, and cytotoxic T cells (CTLs), an important subset of T cells responsible for killing cancer cells.

## Results

### Fabrication of HICs

To model solid tumors, hypoxia-inducing scaffolds were fabricated by cryogelation at −20°C as previously described (Figure 1A).(*29, 30*) Since HA is an important component of the tumor-associated ECM(*36*) that promotes tumor progression,(*37*) polymerizable HAGM (i.e., HA modified with glycidyl methacrylate) was used as the cryogel building block. Integrins binding to ECM ligands are a dominant factor of cancer aggressiveness,(*38*) therefore, chemically modified RGD (APR: acrylate-PEG-G_4_RGDSP),(*39*) a short peptide present in various proteins of the ECM, was copolymerized with HAGM. To induce hypoxia, acrylate-PEG-glucose oxidase (APG) and acrylate-PEG-catalase (APC) were also used as co-monomers to enzymatically deplete O_2_ and degrade H_2_O_2_ byproduct, respectively. After cryo-polymerization (step 1), HICs were thawed at RT to melt ice crystals, resulting in a scaffold with large and interconnected pores. This unique macroporous structure facilitates cell seeding and infiltration throughout the scaffold and allows 3D reorganization of cells to mimic the tumor architecture (step 2). In addition, this porous construct facilitates gas and solute diffusion, improving O_2_ depletion by APG while preventing H_2_O_2_ accumulation. As illustrated in Figure 1B, HICs were engineered to recreate aspects of aggressive solid tumors by emulating (*i*) hypoxia-associated tumor cell metabolism, phenotype, and resistance to chemotherapeutics, (*ii*) hypoxia-associated immune cell inhibition (e.g., dendritic cells), and (*iii*) an immunosuppressive TME, a condition known to induce resistance of tumor cells to CTL-mediated cytotoxicity.

**Figure 1:**
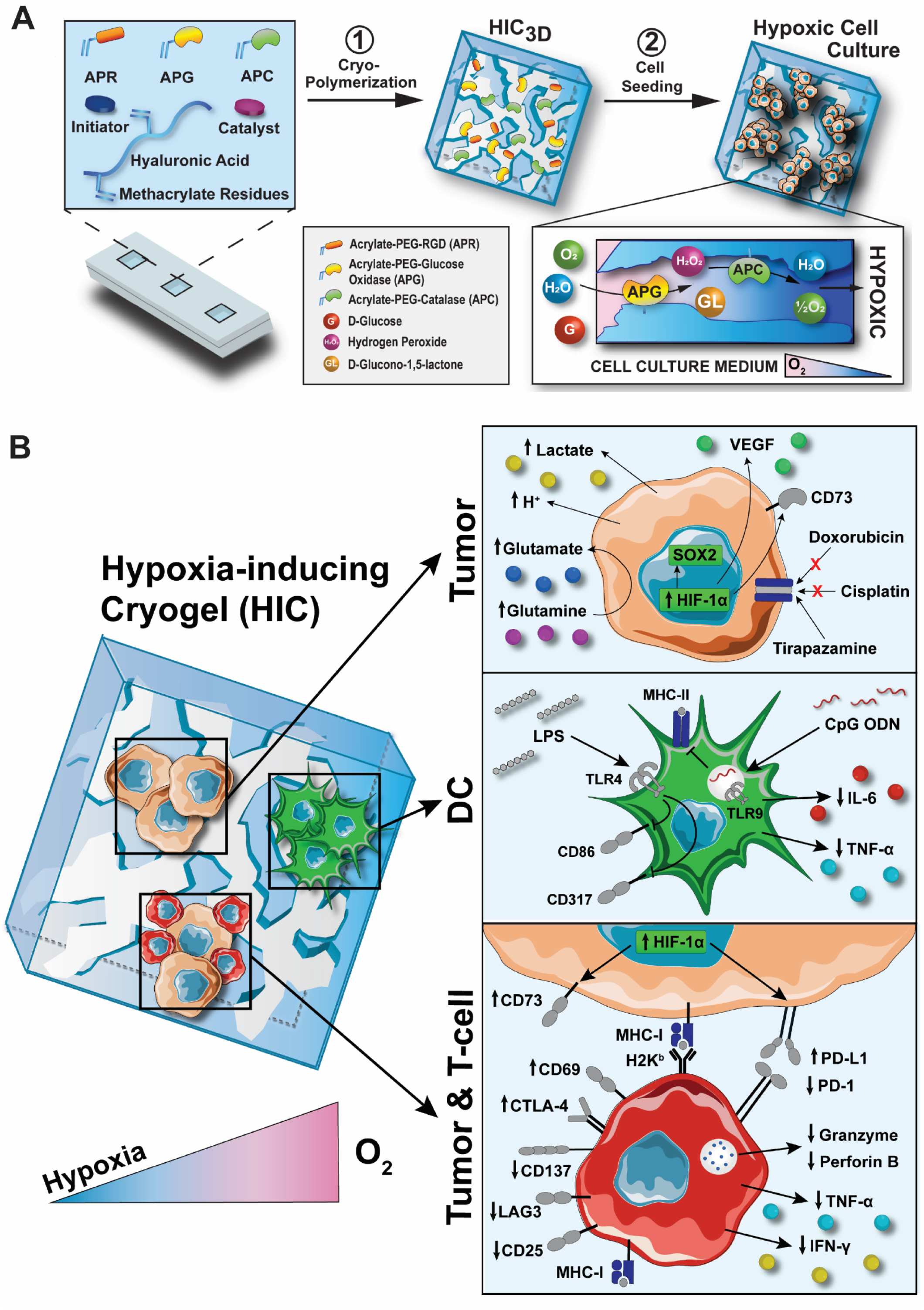
Schematic illustration of the fabrication process of HICs and their ability to induce a hypoxic tumor microenvironment. **A**) HICs are first fabricated by mixing HAGM, APG, APC, APR, and the initiator system (TEMED/APS) in water (H_2_O) and subsequently reacted at subzero temperature (−20°C) for 16h to allow cryogelation (step 1). Once the cryopolymerization is complete, HICs are thawed at RT, washed thoroughly with PBS, transferred into a well plate (1 cryogel/well), seeded with cancer cells or immune cells, and subsequently incubated under normoxic conditions (i.e., within a standard incubator operating at 18.6% O_2_) (step 2). When HICs are immersed in a cell culture medium supplemented with D-glucose, APG depletes O_2_ by catalyzing the oxidation of D-glucose (G) to D-glucono-1,5-lactone (GL) and hydrogen peroxide (H_2_O_2_). The resulting H_2_O_2_ is sequentially hydrolyzed by APC to H_2_O and ½ O_2_, inducing hypoxic conditions (0.5–2% O_2_). **B**) HICs recreate a hypoxic tumor microenvironment and can be leveraged as a tool to investigate the effect of hypoxia on cancer cells such as adaptive changes of their metabolism, phenotype, and response to chemotherapeutics (upper panel). HICs can also be used to induce hypoxia-mediated DC inhibition and phenotypic changes (middle panel) and as a platform to mimic cancer-immune cell interactions in 3D and better understand hypoxia-driven immunosuppression on tumor-reactive T cells (lower panel).

### Characterization of HICs

Cryogels possess unique physical properties including a well-controlled porous structure and mechanical strength resulting in an elastic construct that facilitates biomolecule diffusion, waste removal, and cell infiltration, invasion, and trafficking. To understand the impact of enzyme grafting on cryogels, we first evaluated their microstructure as a function of enzyme concentration. HICs were fabricated with 0.4% APC and various APG concentrations (0.01–0.1%). Blank cryogel (APC-free and APG-free) was used as a control. As shown in Figure 2A-C, confocal microscopy and SEM imaging showed that HICs exhibit large pores, ranging from 20 to 200 µm in diameter. Additionally, no significant changes were observed in pore size across the different APG concentrations investigated as the average pore diameters of HICs and blank cryogels were found to be similar around 50 µm. Similarly, grafting chemically-modified enzymes did not significantly impact cryogels as the pore connectivity (83 ± 3 %) and swelling ratio (Qm = 46 ± 4) were similar across all the groups (Figure 2D-E). However, it is worth noting that the addition of APC and APG slightly reduced the Young’s modulus of cryogels independently of their concentrations, with values of 2.7 ± 0.2 kPa and ∼1.9 ± 0.1 kPa for blank cryogels and HICs, respectively (Figure 2F). Next, to confirm the successful incorporation of the enzymes into the cryogels, HICs were fabricated with fluorescently-labeled APC (fluorescein, green) and APG (cyanine-5, magenta) and analyzed by confocal microscopy (Figure 2G). As expected, the grafting efficiency of APG into the polymer walls was proportional to the concentration used. Importantly, changes in APG concentration did not influence APC incorporation into the cryogel. Finally, the syringe injectability of HICs was tested (Figure 2H, Video 1). All cryogels withstood the syringe injection through a 16G hypodermic needle without any mechanical damage and displayed shape memory properties. Collectively, these results indicate that despite a slight change in mechanical properties compared to blank cryogels, HICs retained the unique and advantageous characteristics of cryogels.

**Figure 2:**
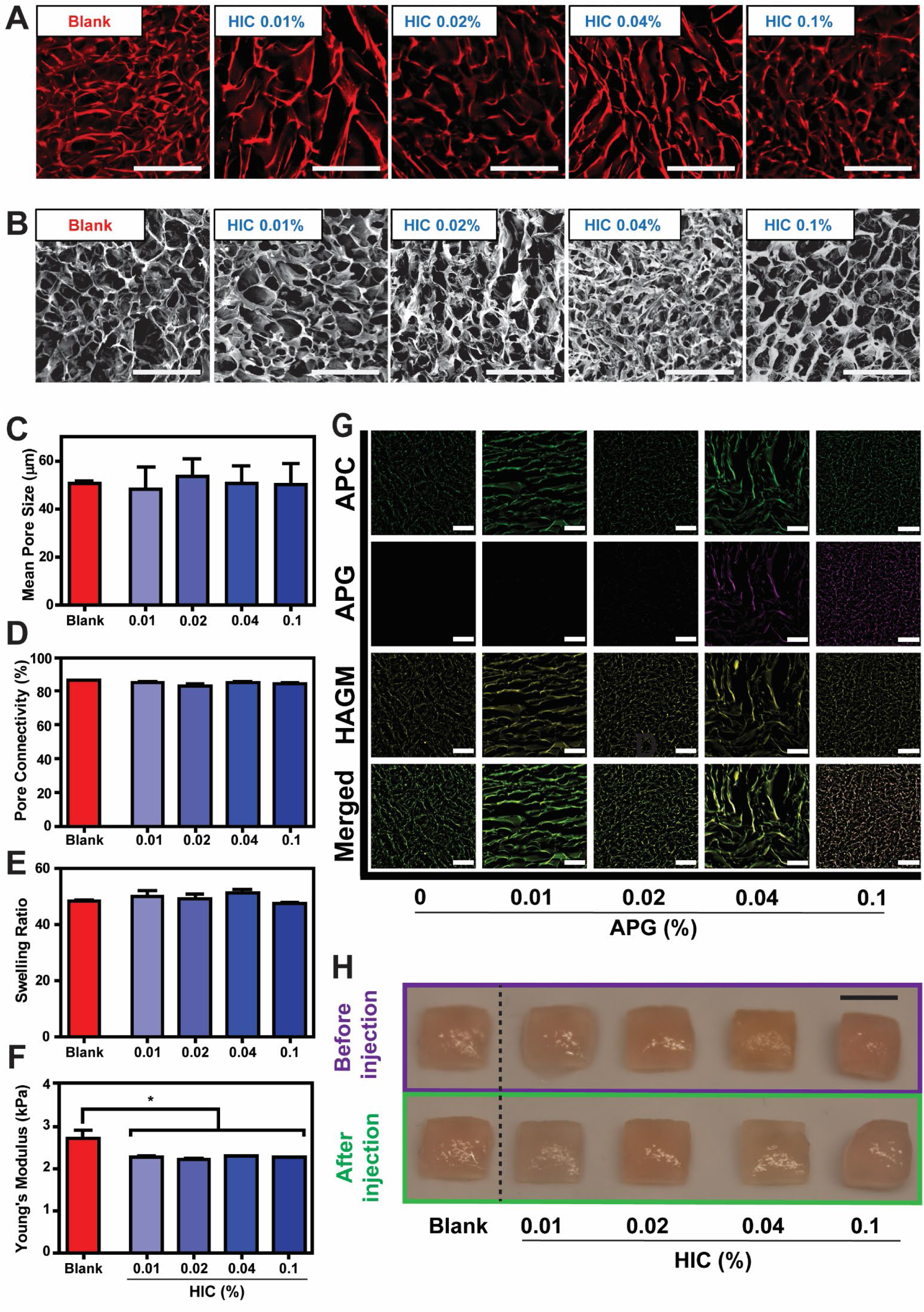
Physical characteristics of HICs. HICs were formulated with APG (0.01, 0.02, 0.04, or 0.1%) and APC (0.4%). Blank cryogels (i.e., APC-free and APG-free) were used as control. **A–B**) Confocal microscopy (A) and SEM (B) images depicting the macroporous and interconnected network of cryogels (blank and HICs). **C–F**) Physical characterization of blank cryogels and HICs: (C) Mean pore size, (D) Pore connectivity, (E) Swelling ratio (Qm), and (F) Young’s modulus. **G**) Confocal images displaying the incorporation efficiency of enzymes (APC and APG) into cryogels. APC was fluorescently labeled with fluorescein (green) and APG with cyanine-5 (magenta). **H**) Photographs of blank cryogels and HICs pre and post-syringe injection. Each image or photograph is representative of n = 5 samples. Values represent the mean ± SEM (n = 5–8 cryogels), and data were analyzed using ANOVA and Dunnett’s post-hoc test (compared to blank cryogels). *p < 0.05. Scale bars = 100 µm (A, B, G) and 4 mm (H).

### Hypoxia characterization and biological studies

We first investigated the capability of HICs to deplete O_2_ in cell culture conditions using contactless O_2_ sensor spots (Figures 3A-B and S1A). In an aqueous solution supplemented with glucose, APG quickly converts D-glucose (C_6_H_12_O_6_) and O_2_ into H_2_O_2_ and D-glucono-1,5-lactone (C_6_H_10_O_6_) (Equation 1). Then, APC comes into play to catalyze the degradation of H_2_O_2_ into H_2_O + ½ O_2_ to prevent H_2_O_2_-related cytotoxicity at concentrations >10 μM (Equation 2).(*40*)

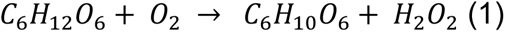

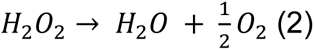

**Figure 3:**
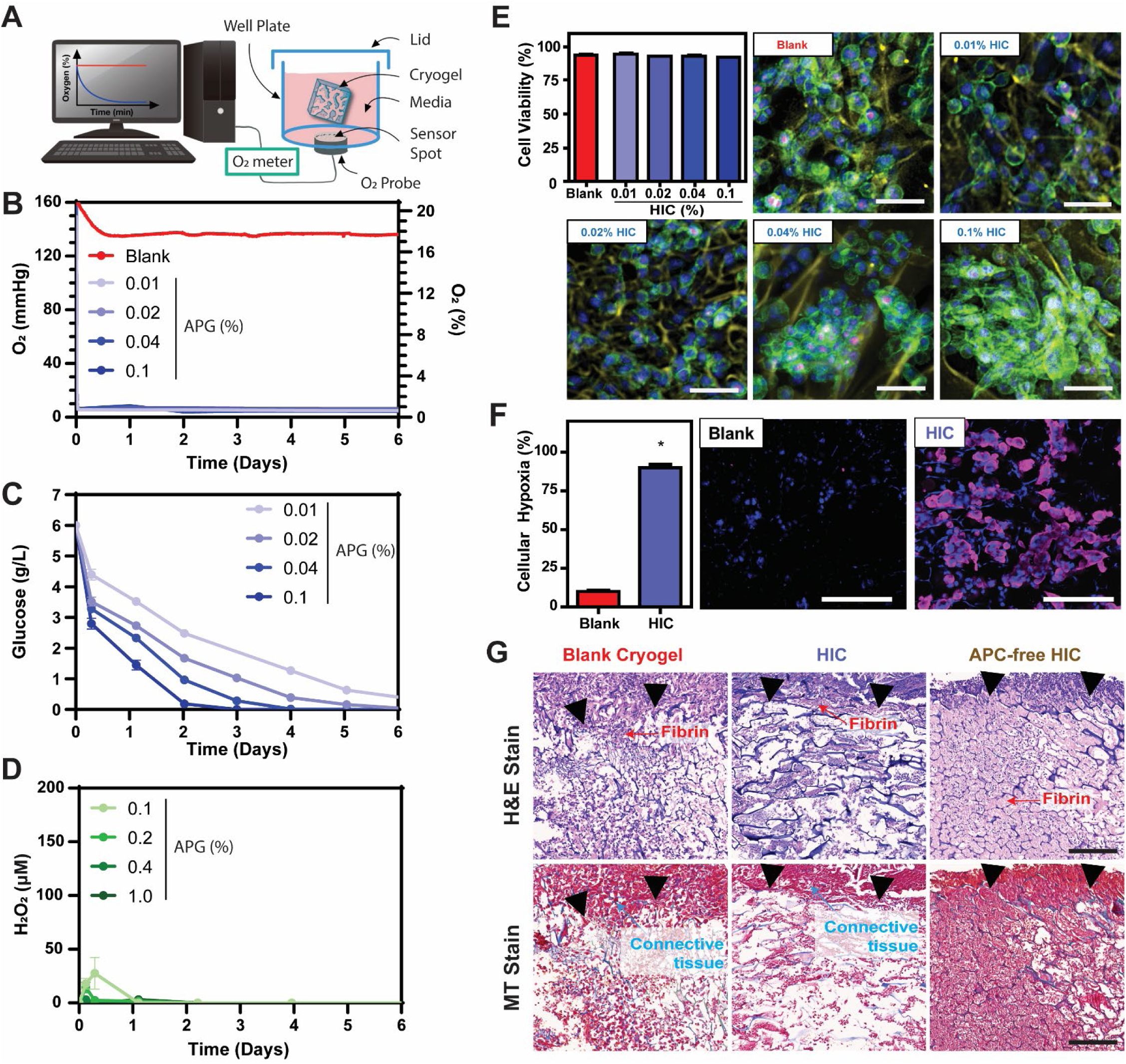
HICs create a hypoxic environment while remaining cytocompatible and biocompatible. **A**) Schematic describing the experimental setup for O_2_ measurements. Blank cryogels or HICs (0.04% APG) were incubated in a p48 well plate at 37°C under normoxic conditions (18.6% O_2_ = 140 mmHg). Each well contained 0.5 mL of complete DMEM and a contactless O_2_ sensor spot was attached to its bottom. Changes in dissolved O_2_ tension were recorded every 30s. **B**) Sustained O_2_ depletion from blank cryogels or HICs. **C–D**) Kinetics of glucose consumption (C) and H_2_O_2_ production (D) from enzymatically active HICs. **E**) Viability and confocal images of B16-F10 cells cultured in blank cryogels or HICs (0.4% APC, 0.01–0.1% APG) after 24 h of incubation. Blue = nuclei stained with DAPI, red = dead cells labeled with ViaQuant Far Red, green = actin cytoskeleton labeled with Alexa Fluor 488 phalloidin, yellow = polymer walls labeled with rhodamine. **F**) Quantification and confocal images of cellular hypoxia of B16-F10 cells cultured in blank cryogels or HICs (0.04% APG, 0.4% APC) after 24 h of incubation. Blue staining = nuclei stained with DAPI, purple = hypoxic cells stained with Hypoxyprobe-1. **G**) Histological analysis. Masson’s trichrome (MT) and hematoxylin and eosin (H&E) staining of cuboidal cryogels (dimensions: 4 × 4 × 1 mm^3^) explanted 28 days following subcutaneous injections in the dorsal flanks of C57BL/6 mice. Samples: blank cryogel (left column), HIC: 0.04% APG + 0.4% APC (middle), and APC-free HIC: 0.04% APG (right column). H&E staining highlights the macroporous polymeric network of cryogels (interconnected dark blue fibers), infiltrated leukocytes (dark blue dots), fibrin formation (purple), and surrounding tissues (cryogel-free). Black arrows indicate the boundary between the cryogel and the host tissue. Confocal and histological images are representative of n = 5 samples per condition. Values represent the mean ± SEM and data (panel F) were analyzed using ANOVA and Dunnett’s post-hoc test (compared to blank cryogels, n = 5). *p < 0.05. Scale bars = 100 µm (E, F) and 100 µm (G).

As expected, HICs incubated in glucose-free DMEM under normoxic conditions did not change the O_2_ tension (Figure S1A). However, in a glucose-supplemented DMEM, HICs induced a rapid depletion of O_2_ across all APG concentrations (0.01–0.1%) tested (Figure 3B), with a time to hypoxia of 2 h, and hypoxic conditions sustained for up to 6 days (Figure S1F-G). Next, kinetics of glucose consumption by HICs were evaluated. HIC-induced glucose depletion was proportional to the amount of APG (0.01–0.1%) grafted into the cryogels with consumption rates ranging from 5.5 ± 0.6 to 10.97 ± 0.6 g/L/day during the hypoxia induction phase (Figures 3C and S1B). Once O_2_ steady state (e.g., hypoxia) was reached, the glucose consumption rates of HICs (0.01–0.1%) significantly decreased, varying from 0.73 ± 0.2 to 1.4 ± 0.6 g/L/day (Figure S1C). Next, the influence of APC concentration on net H_2_O_2_ production was explored. HICs containing 0.04% APG were grafted with various amounts of APC (0.01–0.1%) and subsequently incubated in glucose-supplemented Dulbecco’s phosphate buffered saline (DPBS) for 6 days (Figures 3D, S1D, and S2). As expected, the addition of APC into the cryogels did not impact APG grafting efficiency (Figure S2). HICs containing 0.4 % APC induced low H_2_O_2_ accumulation with a maximum of 3 ± 3 μM during the induction phase, while gels with 1% APC completely prevented net H_2_O_2_ production (Figures 3D and S1D). However, lower amounts of APC led to higher levels of H_2_O_2_ during the initial O_2_ depletion phase, ranging from 14.5 ± 4.9 μM after 3 h to 27.5 ± 14.7 μM after 7 h for HICs with 0.2 and 0.1 % APC, respectively. Interestingly, there was no detection of H_2_O_2_ for all HIC formulations during the O_2_ steady state phase. Furthermore, we investigated the cell viability within HICs as an indication of cytocompatibility. Blank cryogels and HICs were seeded with B16-F10 cells (1 x 10^5^ cells/cryogel) and incubated for 24 h in normoxia (Figure 3E). All B16-F10 cell-laden HICs displayed a homogenous cell distribution throughout the scaffolds with high viability (∼96%), similar to their blank cryogels counterparts. It is worth noting that cell clusters were visible after 24 h within HICs containing the highest amounts of APC (0.04 and 0.1 %), suggesting the capacity of these constructs to promote tumor cell reorganization into 3D-like microtissues. Taken together, HICs containing 0.04 and 0.4% APC were selected and used for the rest of our studies. These HICs have the ability to induce a steady state of hypoxia (∼0.8% O_2_) within 2 h that can last for at least 6 days (Figure S1E-G).

Next, HICs were tested for their capacity to induce cellular hypoxia. B16-F10 cells (1 x 10^5^ cells/cryogel) were cultured in blank cryogels and HICs under normoxic conditions for 24 h, and pimonidazole (detection < 1.5% O_2_) staining was used to assess cellular hypoxia (Figure 3F). After incubation, 90.5 ± 0.8% of the cells within HICs were hypoxic, whereas only 10.3 ± 0.5 % of hypoxic cells were observed in blank cryogels, suggesting an efficient induction of cellular hypoxia by HICs. Finally, the biocompatibility of HICs and APC-free HICs was investigated in mice. Cryogels were first subcutaneously injected in the dorsal flanks of C57BL/6J mice with glucose-supplemented DPBS, then explanted with the surrounding tissues after 3 and 28 days (Figures 3G and S3), and finally analyzed by histology using H&E and MT staining. All excised cryogels retained their initial size and shape. Only a few neutrophils were observed in the HICs and blank cryogels, suggesting minimal host immune reaction and inflammation (Figures 3G and S3). In addition, both types of cryogels were well integrated within the native tissues, as demonstrated by the thin fibrin layer at the junction between the connective tissue stroma and the constructs. In contrast, APC-free HICs showed a high neutrophil infiltration as early as 3 days post-injection and lasting for at least 28 days. In addition, a high fibrin deposition deep inside the constructs was observed with poor integration with the surrounding tissues, indicating a strong and chronic inflammatory reaction. Collectively, these results suggest that HICs can induce cellular hypoxia while being biocompatible. In addition, our data indicate that catalase is key in preventing the induction of a host inflammatory response by residual HIC-induced H_2_O_2_.

### HIC-based cancer models recapitulate tumor hypoxia and resistance to chemotherapy

Several studies have reported that the hypoxic TME has an impact on cancer cell metabolism and phenotype.(*41, 42*) Thus, we investigated the capacity of HIC-supported tumor models to recapitulate the impact of hypoxic conditions on cancer cells. B16-F10 cells were seeded within blank cryogels and HICs (1 x 10^5^ cells/cryogel) and cultured for up to 72 h. A change in the metabolic activity was evaluated by measuring the extracellular pH and lactate concentration, as well as the glutamine and glutamate concentrations as markers of aerobic glycolysis and glutaminolysis, respectively.(*43, 44*) Although pH values remained within the physiological range for cancer cells, HICs induced a notable decrease of extracellular pH compared to blank cryogels: 6.6 ± 0.1 vs. 7.2 ± 0.1 at 72 h, respectively (Figure 4A). In addition, compared to blank cryogels, HICs substantially increased lactate and glutamate secretion (∼3-fold) and decreased glutamine concentration (∼1.5-fold) in the cell culture supernatant (Figure 4B). Next, we investigated the impact of HICs on B16-F10 cell phenotype. Total mRNA was extracted from the cells cultured for 24 and 48 h, and the expression profile of several hypoxia-associated tumor aggressiveness markers was analyzed by RT-qPCR (Figure 4C): *i*) HIF-1α, a subunit of the transcription factor hypoxia-inducible factor 1 implicated in the regulation of cellular and developmental response to hypoxia;(*45*) *ii*) cluster of differentiation 73 (CD73), a surface enzyme responsible for the extracellular accumulation of immunosuppressive adenosine;(*46*) *iii*) vascular endothelial growth factor alpha (VEGFα), a major angiogenic factor that has been associated with cancer progression and decreased survival rates;(*47*) and *iv*) SRY (sex determining region Y)-box 2 (SOX2), an embryonic stem cell transcription factor involved in cancer cell invasion.(*48*) After 48 h of incubation, B16-F10 cells within HICs had a significant increase of HIF-1α (2-fold), VEGFα (12-fold), CD73 (2-fold), and SOX2 (8-fold) compared to the expression levels of B16-F10 cells within blank cryogels. It should be noted that the expression levels of CD73 and SOX2 started to increase just after 24 h of culture within HICs. Surprisingly, B16-F10 cells in blank cryogels also demonstrated a slight overexpression of HIF-1α, VEGFα, and SOX2, most likely due to partial hypoxia within the construct owing to an increased consumption of oxygen relative to cell proliferation. Collectively, these results suggest that HICs emulate the hypoxic TME, inducing hypoxia-mediated metabolism reprogramming and phenotypic changes in cancer cells.

**Figure 4:**
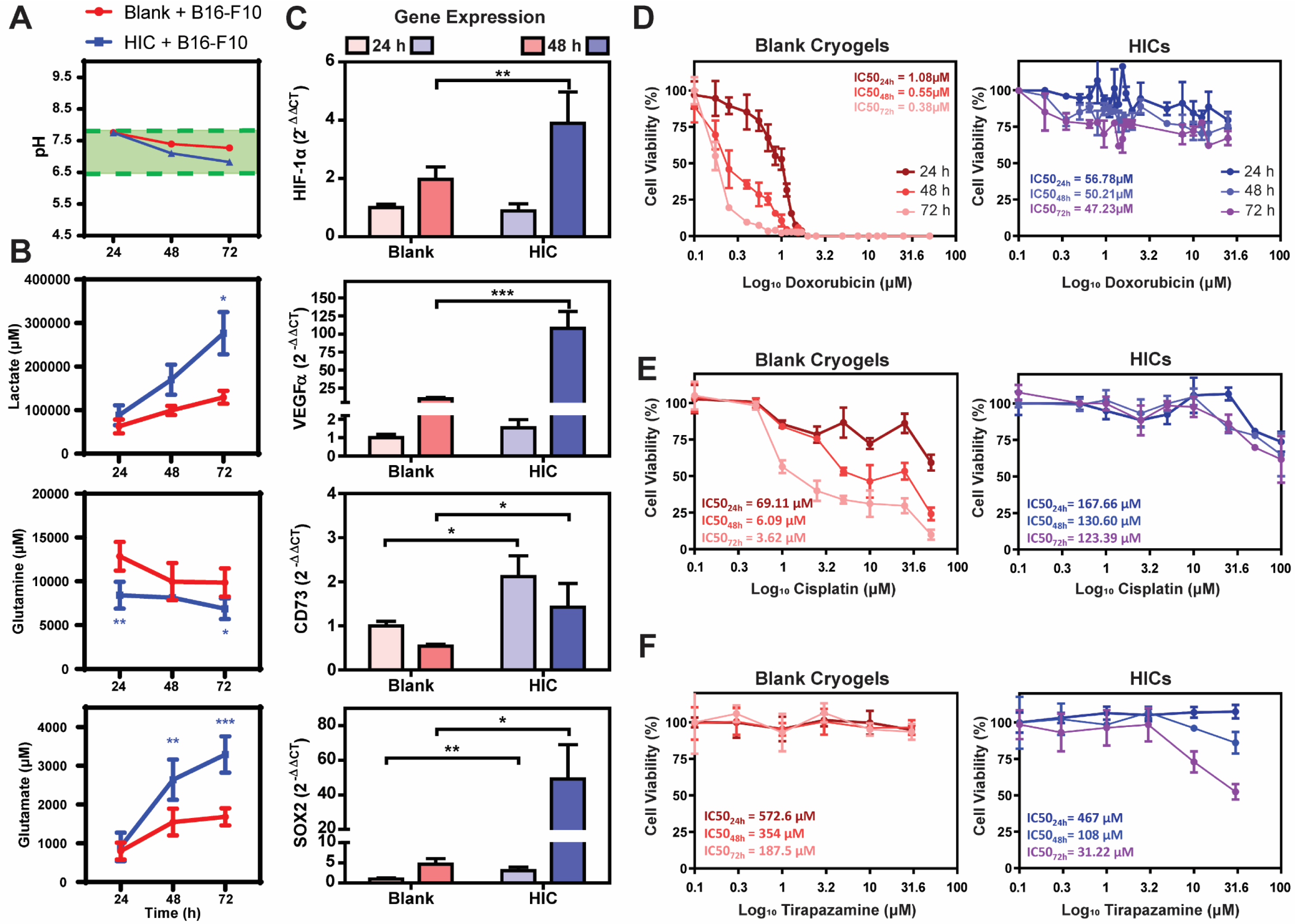
HIC-based cancer models induce hypoxia-induced cell metabolism, phenotype, and resistance to chemotherapeutics. **A**) Quantification of pH change in cell culture medium after 24, 48, and 72 h of incubation of B16-F10 cells cultured in blank cryogels or HICs. The green area represents the physiological pH values of the cell culture media. **B**) Concentrations of Lactate, Glutamine, and Glutamate from the cell culture media of B16-F10 cells cultured in blank cryogels or HICs after 24, 48, and 72 h of incubation. **C**) Gene expression levels of HIF-1α, VEGFα, CD73, and SOX2 in B16-F10 cells cultured in blank cryogels or HICs for 24 and 48 h. **D–F**) Viability of B16-F10 cells cultured in blank cryogels or HICs and treated with doxorubicin (D), cisplatin (E), and tirapazamine (F) at various concentrations (0–31.6 µM) for 24, 48, and 72 h. All cell culture studies were conducted under normoxic conditions (i.e., within a standard incubator operating at 18.6% O_2_). Values represent the mean ± SEM and data were analyzed using ANOVA and Dunnett’s post-hoc test (compared to blank cryogels, n = 5–6). *p < 0.05, **p < 0.01, ***p < 0.001.

Next, we evaluated HIC-induced cancer cell resistance to chemotherapeutic agents. B16-10 and 4T1 cells (1 x 10^5^ cells/per cryogel) were seeded within blank cryogels and HICs, treated with doxorubicin (DOX) and cisplatin (CIS) at various concentrations (0–31.6 μM), and cell viability was assessed after 24, 48, and 72 h of incubation (Figures 4D-E, S4A-B, and S5A-B). The half-maximal inhibitory concentration (IC_50_) was determined as the dose of chemotherapeutic agents killing 50% of cancer cells. Tirapazamine (TPZ), an anti-cancer drug activated only in hypoxic conditions, was used as a control. Our data show that B16-F10 cells cultured in blank cryogels were extremely sensitive to DOX (Figures 4D and S4A) and CIS (Figure 4E) treatments, with IC_50_ values decreasing from 1.08 μM and 69.11 μM after 24 h to 0.38 μM and 3.62 μM after 72 h, respectively. In contrast, B16-F10 cells in HICs demonstrated high resistance to both chemotherapeutics after 72 h of treatment, with IC_50_ values of 47.23 μM for DOX and 123.39 μM for CIS, which represents an increase in chemoresistance by 124-fold and 34-fold, respectively, compared to our controls (i.e., cells cultured in blanks cryogels). Similar results were obtained with 4T1 cells, with an increased resistance to DOX by 290-fold and to CIS by 101-fold in comparison to our controls (Figures S4B and S5A-B). As expected, TPZ had little impact on the viability of B16-F10 cells in blank cryogels (IC_50_ = 187.5 μM), while inducing significant cell mortality in those cultured within HICs (IC_50_ = 31.22 μM). Overall, these results indicate that HICs can locally emulate a hypoxic TME, bestowing cancer cells hypoxia-mediated resistance to chemotherapy drugs or sensitivity to hypoxia-activated pro-drugs.

### HICs induce DC subset change and immunomodulation

Next, we tested the capacity of HICs to mimic hypoxia-mediated inhibition in a tumor-like microenvironment. Hypoxia is a key microenvironmental factor in solid tumors that inhibits immune cells. It can negatively impact DC survival and activation,(*49, 50*) as well as impair the stimulation and cytotoxic activity of tumor-reactive T cells.(*51*) However, some studies also reported divergent effects of hypoxia on DCs such as increased expression of co-stimulatory molecules or a potentializing effect upon stimulation with adjuvants.(*52–54*) Therefore, we first investigated the effect of hypoxic conditions on primary DC function (Figures 5A and S6A). Bone marrow-derived dendritic cells (BMDCs) were stimulated for 24 h with lipopolysaccharide (LPS) (Figure 5) or cytidine-guanosine-dinucleotides oligodeoxynucleotide (CpG ODN) 1826 (Figure S6) in normoxic or hypoxic conditions in 2D (i.e., cryogel-free) or 3D (i.e., blank cryogels). Although no changes in DC phenotype were observed in the non-treated conditions, LPS and CpG-mediated stimulation significantly decreased the fractions of plasmacytoid B220^+^ DCs over conventional B220^−^ DCs in normoxia (Figures 5B and S6B). Conversely, in hypoxia, both stimulations induced an increase in the B220^+^ DC population in 2D compared to blank cryogels (60% *vs.* 30%). Interestingly, B220^+^ and B220^−^ DC subsets displayed different activation and maturation markers in hypoxia upon LPS and CpG stimulation (Figures 5C-D and S6C-D). LPS-treated B220^+^ DCs in 2D exhibited an increased expression of MHCII (∼40%) and CD317 (∼30%) but a decreased expression of CD86 (∼40%) in hypoxia compared to normoxia, whereas a diametrically opposite response was observed with B220^−^ DCs (Figure 5C-D). Similar results were observed with CpG stimulation (Figure S6C-D). Intriguingly, DCs stimulated in the presence of blank cryogels displayed an opposite response compared to 2D. For instance, while the fraction of B220^+^ CD86^+^ DCs decreased upon LPS stimulation in 2D hypoxia *vs.* normoxia (∼15% and ∼60%, respectively), it increased with blank cryogels (∼30% and 10%, respectively). Similarly, the fraction of B220^−^ CD86^+^ DCs was increased in 2D under hypoxic conditions following LPS stimulation compared to normoxia (∼90% *vs*. ∼40%, respectively), whereas it was reduced with blank cryogels (∼50% *vs.* 90%, respectively).

**Figure 5:**
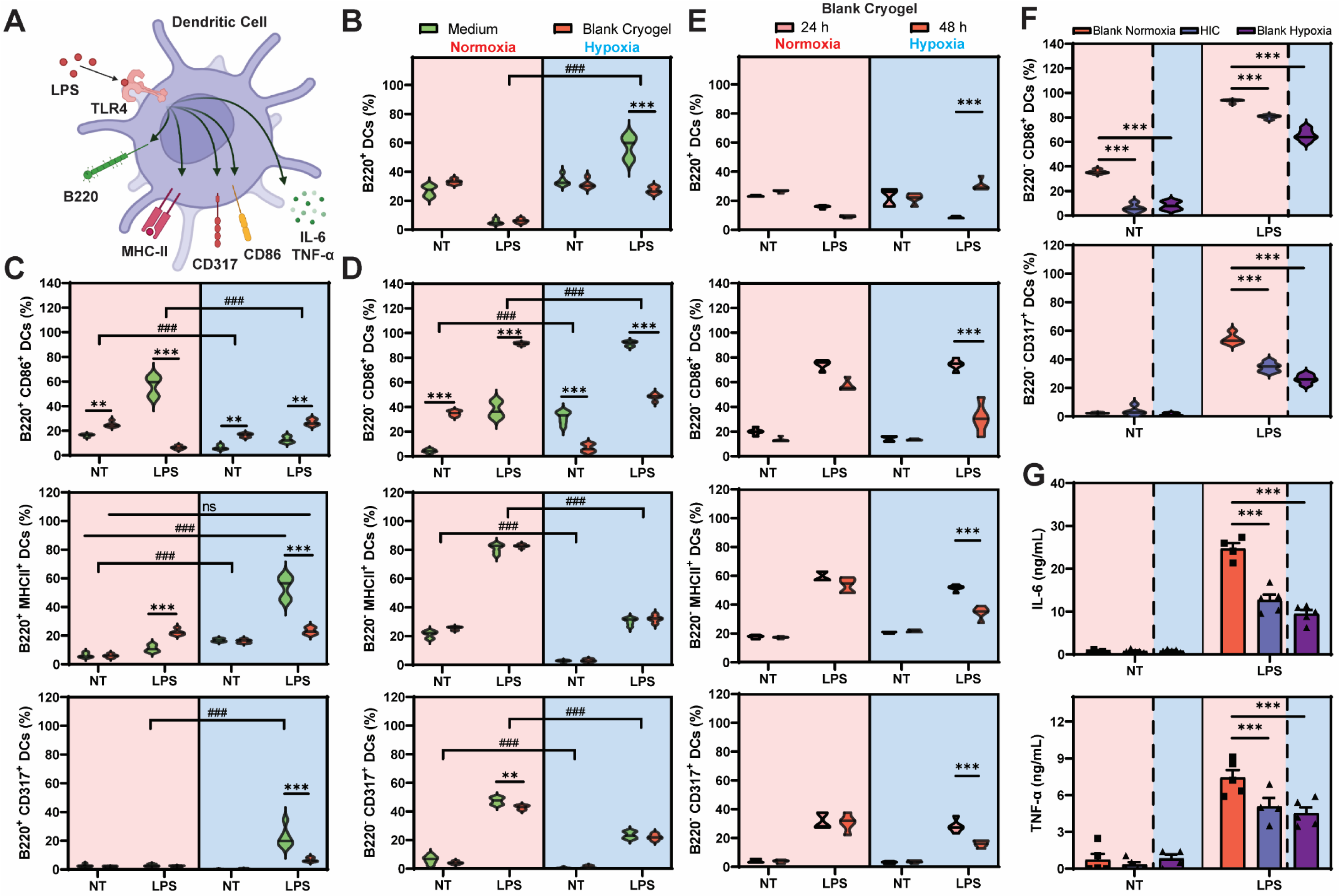
HICs influence subset change and LPS-induced activation of DCs. **A**) Schematic depicting LPS-induced DC activation via TLR4 stimulation under normoxic conditions, leading to increased expression of specific cell surface markers of maturation and production of proinflammatory cytokine. **B**) Fractions of B220^+^ CD11b^+^ CD11c^+^ DCs after 48 h of incubation in cryogel-free and blank cryogel-containing cell culture media under normoxic or hypoxic conditions. DCs were first pre-incubated for 24 h and subsequently treated with LPS for another 24 h. **C–D**) Fractions of activated B220^+^ (C) and B220^−^ (D) DCs associated with increased expression of CD86, MHCII, or CD317. **E**) Fractions of B220+ CD11b+ CD11c+ DCs and activated B220-DCs cultured in blank cryogel-containing cell culture media without (24 h) or with (48 h) pre-incubation under normoxic or hypoxic conditions. **F–G**) Evaluation of DC activation (F) and cytokine secretion (G) in the presence of blank cryogels or HICs. (F) Fractions of activated DCs associated with increased expression of CD86 or CD317. (G) Proinflammatory cytokine secretion. Concentrations of IL-6 and TNF-α in cell supernatants were measured by ELISA. Across all studies, non-treated DCs (NT) were cultured in LPS-free medium and used as control. Cell culture studies were conducted under normoxic (i.e., within a standard incubator operating at 18.6% O_2_) or hypoxic (i.e., within a tri-gas incubator operating at 1% O_2_) conditions. Values represent the mean ± SEM (n = 5) and data were analyzed using ANOVA and Dunnett’s post-hoc test. **p < 0.01, ***p < 0.001, ^###^p < 0.001 (*compared to cell culture media or blank cryogels, ^#^compared to normoxia).

Recent studies have shown that HA can induce immunostimulation or immunomodulation based on its molecular weight,(*55, 56*) but also how it interacts with immune cells.(*57, 58*) To expand our previous analysis, we investigated how HA influenced DC subsets and activation states in normoxia and hypoxia. DCs were stimulated with LPS or CpG, in the presence of blank cryogels (composed of chemically modified HA), PEGDM cryogels (neutral polymer: 3D control), or in 2D (wells coated with HA, positive control) (Figure S7A). As anticipated, not only stimulated DCs behaved similarly in blank cryogels and HA-coated conditions but opposite results were obtained with polyethylene glycol dimethacrylates (PEGDM) cryogels as previously seen with the cryogel-free condition (Figures 5B-D and S6B-D). In normoxia, higher fractions of B220^−^ CD86^+^ and B220^−^ MHCII^+^ DCs were observed in HA-containing conditions compared to PEGDM cryogels. On the contrary, in hypoxic conditions, PEGDM cryogels induced an increase of the B220^+^ DC subset and B220^−^ CD86^+^ fraction compared to blank cryogels and HA-coated wells. Altogether, our results first suggest that HA impacts DC differentiation in hypoxia, preventing phenotypic switching from conventional to plasmacytoid. In addition, our data indicate that HA also changes the responses of LPS and CpG-treated DCs within the same subsets, with strong inhibition when subjected to hypoxia.

Another important consideration is the exposure time to hypoxia, as previously observed with cancer cells (Figure 4). Therefore, we investigated how hypoxia pre-conditioning influenced DC activation and maturation. Unconditioned and pre-conditioned DCs for 24 h in normoxia or hypoxia were stimulated with LPS (Figure 5E) or CpG (Figure S6E) for 24 h in the presence of blank cryogels. Unconditioned and pre-conditioned DCs in normoxia did not show any differences in B220^+^ subset fractions or the activation and maturation levels (B220^−^ CD86^+^, CD317^+^, and MHCII^+^). However, in hypoxic conditions, unconditioned DCs displayed the same patterns as normoxic DCs with a lower B220^+^ fraction and resistance to hypoxia-mediated immunosuppression in comparison to pre-conditioned DCs. Finally, we investigated whether HICs could recapitulate hypoxia-driven DC inhibition. BMDCs were first pre-incubated with blank cryogels or HICs and in hypoxia and with blank cryogels in hypoxia. Next, BMDCs were treated with LPS (Figures 5F and S7B) or CpG (Figure S6F) for 24 h. In normoxia, HICs significantly hampered DC activation with decreased fractions of CD86^+^ (∼15%), CD317^+^ (∼20%), and MHCII^+^ (∼40%) DCs compared to blank cryogels. Strikingly, DC inhibition with HICs was comparable to DCs stimulated in blank cryogels under hypoxic conditions (Figures 5F, S6F, and S7B). This finding was further confirmed by evaluating the concentrations of pro-inflammatory cytokines such as IL-6 and TNF-α in the cell culture supernatant (Figures 5G and 6G). HICs significantly decreased the secretion of IL-6 (∼2-fold) and TNF-α (∼1.5-fold) by DCs stimulated in normoxia compared to blank cryogels, similar to hypoxia-mediated DC inhibition in blank cryogels. Collectively, these data suggest that pre-conditioning in hypoxia is critical to understand hypoxia-mediated immunosuppression of DCs. In addition, our results confirm that HICs induce an immunosuppressive hypoxic environment, altering DC activation and cytokine secretion.

**Figure 6:**
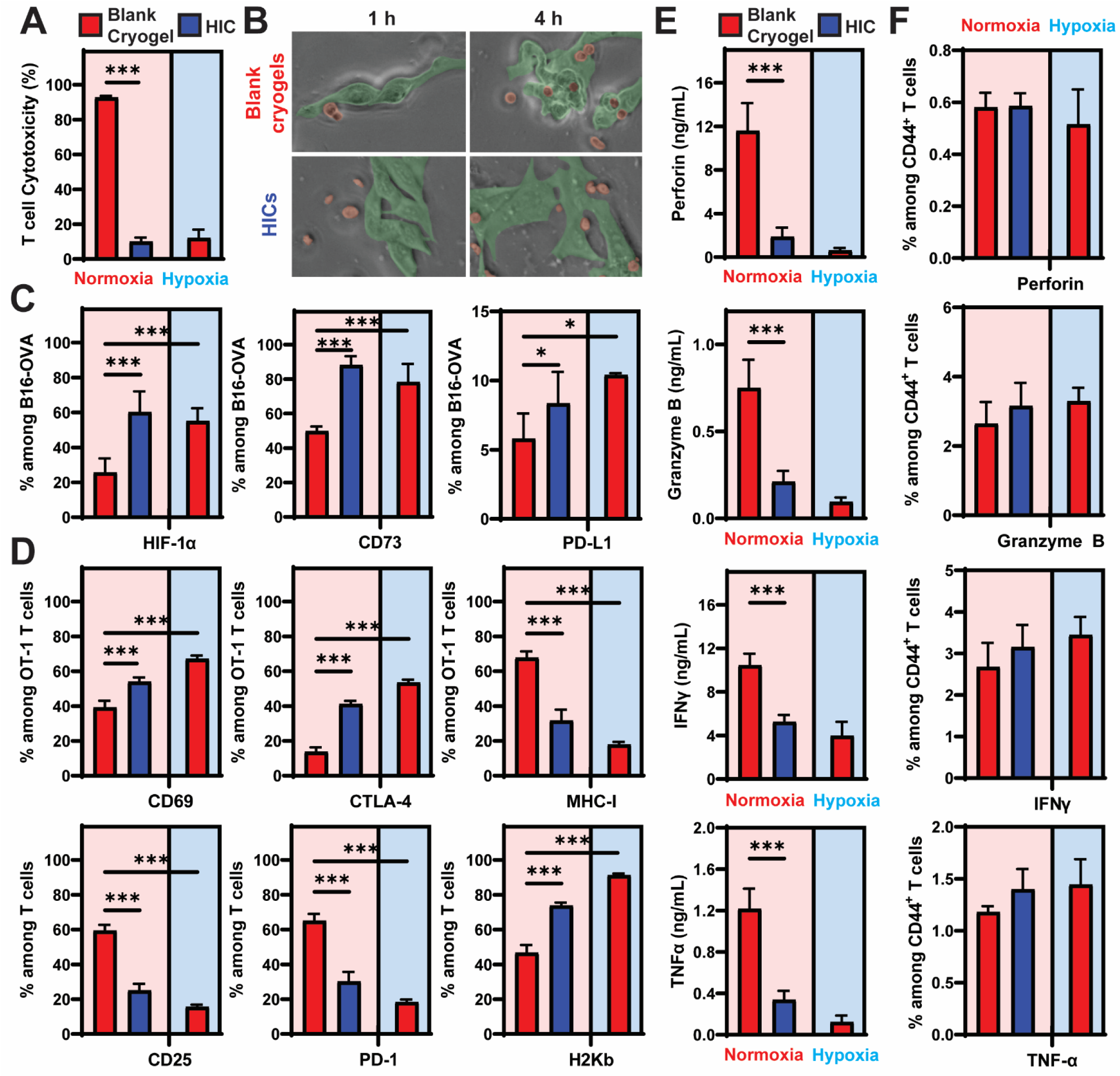
HICs induce hypoxia-driven T-cell immunosuppression. Activation and cytotoxic activity of OT-1 T cells against B16-OVA cells. Cells were co-cultured within blank cryogels or HICs for 24 h under normoxic or hypoxic conditions. **A**) Quantification of OT-1 T cell-mediated cytotoxicity. **B**) Time-lapse images demonstrating OT-1 T cell-mediated attack against B16-OVA in blank cryogels and HICs after 1h (top) and 4h (bottom) of incubation. B16-OVA and OT-1 T cells are pseudocolored in green and red, respectively. **C**) Fractions of B16-OVA cells overexpressing HIF-1α, CD73, and PD-L1. **D**) Fractions of OT-1 T cells overexpressing CD69, CTLA-4, MHC-I, CD25, PD-1, and H2Kb. **E**) Concentrations of secreted perforin, granzyme B, IFNγ, and TNF-α from OT-1 T cells. **F**) Frequencies of cytokine-producing CD44^+^ OT-1 T cells post-stimulation with SIINFEKL peptide. All cell culture studies were conducted under normoxic conditions (i.e., within a standard incubator operating at 18.6% O_2_). Values represent the mean ± SEM (n = 5–6) and data were analyzed using ANOVA and Dunnett’s post-hoc test (compared to blank cryogels in normoxia). *p < 0.05, ***p < 0.001.

### HIC-supported melanoma model recapitulates hypoxia-mediated T cell inhibition

To substantiate our previous findings, we investigated the capacity of HICs to emulate tumor-mediated T cell inhibition in a hypoxic environment. To this end, the HIC platform was used to develop a hypoxic 3D co-culture system in which the cytotoxic function of OT-1 T cells on B16-OVA cells was evaluated (Figure 6A-B, Videos 2,3). B16-OVA cells (1 x 10^5^ cells/per cryogel) were cultured for 3 h within blank cryogels or HICs in normoxia, then subsequently cocultured with OT-1 T cells (ratio 2:1) for another 24 h in normoxia or hypoxia. Under hypoxic conditions, the cytotoxic activity of OT-1 T cells decreased dramatically (∼13 ± 5%), confirming that hypoxia inhibits cytotoxic T cells in this model (Figure 6A). Under normoxic conditions, OT-1 T cells infiltrated the blank cryogels and efficiently killed B16-OVA cells (90 ± 1% cell death) after 24 h. In contrast, HICs protected B16-OVA cells from OT-1 T cells by compromising their cytotoxic activity (∼12 ± 3% cell death), comparable to hypoxia-mediated inhibition (i.e., B16-OVA and OT-1 T cells cocultured in a hypoxic incubator). To further confirm these results, the tumoricidal capacity of OT-1 T cells was assessed by time-lapse microscopy. Under normoxic conditions, B16-OVA cells (1 x 10^5^ cells/well) were cultured for 3 h in 48-well plates in the presence of blank cryogels or HICs, then subsequently cocultured with OT-1 T cells (ratio 2:1) for 4 h (Figure 6B, Videos 2,3). As expected, HICs prevented OT-1 T cells from inducing high tumor cell death, whereas most cells were killed in the presence of blank cryogels.

To better understand how HICs and hypoxia impact OT-1 T cell cytotoxicity, we characterized B16-OVA and OT-1 T cells after 24 h of co-incubation within blank cryogels and HICs in normoxic and hypoxic conditions (Figures 6C-D and S8). The fractions of B16-OVA cells expressing HIF-1α, CD73, and the checkpoint inhibitor programmed death-ligand 1 (PD-L1) significantly increased in HICs and hypoxia (2 to 3-folds) compared to B16-OVA cells cultured in blank cryogels under normoxic conditions (Figure 6C). Surprisingly, HICs did not impact the frequency of CD39^+^ B16-OVA cells, and none of the conditions influenced MHC-I expression on B16-OVA cells (Figure S8A). With respect to OT-1 T cells, we first confirmed that the different coculture conditions did not change the frequency of CD8^+^ OT-1 T cells (Figure S8B). When cocultured in normoxia, we observed that HICs significantly increased the fraction of CD69^+^ (early activation marker), CTLA-4^+^ (checkpoint inhibitor), and H2Kb^+^ (H2Kb ova-specific OT-1 T cell receptor) OT-1 T cells, with frequencies comparable to those obtained under hypoxic conditions (Figure 6D). Similarly, HICs substantially decreased the fraction of MHC-I^+^, CD25^+^ (component of the IL-2 receptor), CD44^+^ and LAG3^+^ (activation markers), CD137^+^ (co-stimulatory receptor), and PD-1^+^ (checkpoint inhibitor) OT-1 T cells. To deepen our understanding, we evaluated cytokine secretion and production by OT-1 T cells after 24 h of co-incubation with B16-OVA cells in normoxia and hypoxia. Under normoxic conditions, compared to blank cryogels, HICs led to a significant decrease in the secretion of perforin (a glycoprotein responsible for pore formation in cell membranes of target cells), granzyme B (a cytotoxic serine protease inducing apoptosis and mediating the cytotoxicity against B16-OVA cells), IFN-γ (mediator of cytotoxic T cell motility and cytotoxicity), and TNF-α (mediator of cytotoxic T cell activation) (Figure 6E). These findings were comparable to hypoxia-mediated inhibition when B16-OVA and OT-1 T cells were cocultured in a hypoxic incubator. Surprisingly, HICs and hypoxia did not alter the ability of OT-1 T cells to produce those biomolecules (Figure 6F), with similar fractions of CD44^+^ OT-1 T cells producing perforin, granzyme B, IFN-γ, and TNF-α compared to blank cryogels in normoxia. Collectively, these results clearly suggest that HIC-supported cancer models induce hypoxia-mediated immunosuppression by interfering with important tumor killing functions of effector T cells.

## Discussion

Hypoxia in the TME is a major contributor to tumor development, aggressiveness, multi-drug resistance, and immune escape.(*5*) Yet, there are no reliable cancer models that mimic both the architecture of tumors and hypoxic conditions.(*59*) In addition, an ideal cancer model should be able to allow immune cell infiltration and recapitulate tumor-immune interactions to advance our knowledge of cancer immunology and improve preclinical screening of immunotherapies.(*60*) To overcome these challenges, we developed an new cryogel-based cancer model platform by creating HICs capable of inducing a local hypoxic tumor microenvironment in 3D while enabling cancer-immune cell interactions.

We engineered a new class of macroporous and syringe-injectable cryogels that can deplete O_2_ in a glucose-dependent and sustained fashion for several days, creating local hypoxic conditions leading to cancer cell-associated metabolic switch, phenotypic changes, and hypoxia-mediated immune cell inhibition. These HICs were fabricated by covalently incorporating APG and APC within the cryogel polymer network, inducing a fast and long-lasting O_2_ depletion while preventing H_2_O_2_-related cytotoxicity. Among all formulations investigated, HICs containing 0.04% APG and 0.4% APC promoted 3D cancer cell reorganization and cellular hypoxia, while consuming the lowest amount of glucose and generating negligible levels of H_2_O_2_. These HICs created a hypoxic environment (∼0.8% O_2_) within 2 h and maintained hypoxia for at least 6 days, whilst remaining cytocompatible and biocompatible. O_2_ depletion could be further prolonged by increasing the amount of APG on cryogels, as well as the amount of APC to prevent any potential cytotoxicity. However, this could lead to unsustainable glucose depletion from the medium that may impact the cell behavior. Other strategies could be investigated to induce hypoxia such as the incorporation of laccase and ferulic acid within cryogels,(*61*) or the encapsulation of O_2_ scavengers like sodium sulfite with the catalyst cobalt sulfate.(*62*)

O_2_-depleting biomaterials have the potential to improve solid tumor modeling *in vitro*, by not only providing adequate biomechanical cues to support cell growth and reorganization in an ECM-like matrix, but also by controlling O_2_ depletion to emulate the hypoxic TME.(*63*) In hypoxia, solid tumors heavily rely on glycolysis over oxidative phosphorylation via the Warburg effect,(*42, 64, 65*) resulting in lactate and glutamate production as well as glutamine uptake and acidosis.(*66*) In this work, we demonstrated that HICs induced hypoxic conditions leading to a pH and glutamine concentration decrease in the cell culture supernatant which was correlated with an increased secretion of glutamate and lactate. This resulted in a profound change in B16-F10 cancer cell metabolism, switching from aerobic respiration to anaerobic glycolysis.(*67*) In addition, hypoxia is a critical factor in cancer aggressiveness and resistance to therapy.(*68*) Here, we showed that HICs promoted the upregulation of HIF-1, VEGFα, and SOX2 genes by B16-F10 cells which have been previously associated with tumor vascularization, as well as tumor cell proliferation, migration, and invasion.(*69–71*) Moreover, cancer cells in HICs displayed similar responses to chemotherapeutic agents used in clinical trials.(*72, 73*) HICs increased the resistance of B16-OVA and 4T1 cells to two commonly used chemotherapeutic drugs to treat cancer, namely DOX and CIS,(*74*) and an increased sensitivity to TPZ, a hypoxia-activated prodrug.(*75*) Collectively, these findings suggest that HICs contribute to hypoxia-mediated metabolism adaptation, phenotypic changes toward aggressiveness, and cancer therapeutic resistance, emulating the behavior of hypoxic solid tumors. However, more work could be performed to optimize HIC-based cancer models, such as investigate the influence of other components of the tumor ECM (e.g., collagen) or matrix stiffness on cancer cell behavior in hypoxia.(*76*) In addition, HIC-based cancer models could be leveraged to investigate the expression of hypoxic biomarkers for cancer therapies,(*77*) as a tool for precision medicine, or to study hypoxia-mediated resistance to other anti-cancer therapies such as radiotherapy.(*78*)

Tumor hypoxia is known to mitigate immune cell infiltration and function in the TME, leading to immune escape.(*79*) However, the *in vivo* exploration of the complex hypoxia-immune cell relationship is challenging, and current hypoxic cancer immunology models are limited.(*80*) In this work, we first demonstrated that several parameters are critical to faithfully emulate *in vitro* DC inhibition observed in the hypoxic TME. We showed that not only does hypoxia influence DC maturation and polarization into pDCs (B220^+^ CD11c^+^) or cDCs (B220^−^ CD11c^+^), but also that HA plays a critical role in these phenotypic changes. In the absence of HA, DCs differentiate in pDCs under hypoxic conditions upon LPS or CpG stimulation. pDCs are a subset of DCs that can be stimulated in hypoxia as demonstrated by the increased fraction of B220^+^ MHCII^+^ and B220^+^ CD317^+^ pDCs. Conversely, in the presence of HA, DCs retained their cDC phenotype in hypoxia, showing a significant alteration in their capacity to be activated by LPS and CpG. This can be correlated with a recent study that reported pDC infiltration in solid triple negative breast cancer as a marker of good prognosis.(*81*) Moreover, we also demonstrated that HA influenced the activation state of DCs within the same subset. pDCs demonstrated an increased B220^+^ CD86^+^ fraction in hypoxia and decreased B220^+^ MHCII^+^ and B220^+^ CD317^+^ fractions in presence of HA. Conversely, while HA increased the B220^−^ CD86^+^ fraction of LPS and CpG-stimulated cDCs in normoxia, it dramatically hampered their activation in hypoxic conditions as compared to HA-free conditions. Therefore, our data indicate that recapitulating the hypoxic tumor immunology in vitro also requires to emulate more closely the TME. This agrees with previous findings demonstrating the crucial role of CD44 (a HA-binding cell surface receptor) in promoting DC activation and DC-T cell immunological synapse.(*82, 83*) Additionally, hypoxia promotes CD44 expression in DCs subjected to hypoxia, leading to a TH2-mediated immune response and cancer immune escape.(*84, 85*) Interestingly, hypoxia-driven inhibition was influenced by the duration of hypoxia prior to stimulation. DC activation was not altered in hypoxic conditions if directly stimulated with LPS or CpG, while a 24-h pre-incubation was sufficient to significantly alter their stimulation. Therefore, at least a 24-h incubation time in 1% O_2_ seems to be necessary to induce hypoxia-mediated DC changes, similar to what was observed with B16-F10. This could explain the divergence in the field on the effect of hypoxia on DC activation(*86, 87*), as a slight change in the incubation time and O_2_ tension could significantly influence immune cell fate. Finally, we demonstrated that HIC-based immune activation models can recapitulate hypoxia-driven immunosuppression. Upon LPS and CpG stimulations, HICs reduced considerably the fractions of activated CD86^+^, CD317^+^, and MHCII^+^ cDCs and the secretion of pro-inflammatory cytokines (IL-6 and TNF-α) to levels comparable to those cultured in our control (blank cryogels) under hypoxic conditions. Overall, we believe the HIC platform is able to engender more closely the behavior of immune cells in solid tumors. However, more studies are required to fully dissect how time to hypoxia and the tumor ECM influence immune cell activation and maturation. For instance, using other ECM molecules (e.g., collagen, fibronectin, elastin, and laminin) as the main components of the cryogel matrix could further give us some insights on how immune cells behave in hypoxia.(*88–91*)

Finally, we demonstrated that HIC-based cancer immunology models could prevent T cell-mediated antitumor activity, with inhibition levels comparable to those observed in hypoxia.(*39*) We observed that HICs and hypoxic conditions markedly increased the fraction of B16-OVA cells overexpressing HIF-1α, as well as CD73 and PD-L1 which are known to play a critical role in inhibiting immune cells via (*i*) the production of the immunosuppressive adenosine(*92, 93*) and (*ii*) the binding to PD-1, a cell surface receptor known to turn off tumor-specific T cells,(*94*) respectively. Similarly, we showed that HICs emulate hypoxia-driven T cell inhibition, with a substantial decrease in the fraction of OT-1 T cells expressing IL-2, CD25, and co-stimulatory TNF receptor CD137, ordinarily responsible for T cell activation and survival.(*95*) This agrees with a decreased secretion of pro-inflammatory cytokines such as IFN-γ and TNF-α observed in the cell culture supernatant of HICs and hypoxic conditions. On the contrary, we showed an increase in the frequency of CD69^+^ OT-1 T cells confirming their early activation,(*96*) and H2Kb^+^ OT-1 T cells, demonstrating high levels of the OVA-specific transgenic T cell receptor that is responsible for B16-OVA recognition.(*97*) Interestingly, although HICs and hypoxic conditions promoted the expression of the inhibitory immune checkpoint CTLA-4 on cytotoxic T cells, we observed a lower fraction of OT-1 T cells expressing the checkpoint inhibitors PD-1 and LAG3.(*46, 98*) Considering that B16-OVA cells overexpressed PD-L1, this suggests that PD-L1/PD-1 interaction is not the primary mechanism responsible for T cell inhibition in our study. Similarly, our results indicate that strategies targeting CTLA-4 may be more promising against solid tumors subjected to hypoxia. Surprisingly, although MHC-I expression was decreased in OT-1 T cells when cultured in HICs or under hypoxic conditions, no changes were observed in B16-OVA cells suggesting that an alteration of MHC-I-H2Kb interaction is not the main driver for inhibiting T cell-mediated cytotoxicity. However, with the decrease of MHC-I, hypoxic T cells may be more susceptible to natural killer (NK) cell-mediated death. Studies have shown that NK cells can modulate T cell-mediated immunity via direct killing in the context of chronic inflammation,(*99*) graft-versus-host diseases,(*100*) and viral infections.(*101*) Therefore, one of the mechanisms of tumor immune escape in hypoxia could be hijacking NK cell function and the killing ability against T cells to reduce the immune system’s ability to recognize, target, and eliminate cancer cells. Furthermore, we clearly quantified a substantial decrease of T cell-mediated secretion of cytotoxic proteins (perforin and granzyme B), which may explain why B16-OVA cell death was prevented in HICs and hypoxic conditions. Surprisingly, we did not observe any change in the production of those cytotoxic molecules and pro-inflammatory cytokines within OT-1 T cells when compared to normoxia. Based on our previous observations(*39*) and these new findings, hypoxia may not be preventing the interactions between cancer cells and T cells but rather creating an immunosuppressive environment in which T cell activation is hindered, halting the release of cytotoxic proteins to give the final blow to cancer cells. This could be attributed to the accumulation of extracellular adenosine(*93*) or other immunosuppressive molecules, or the lack or downregulation of T-cell stimulatory and co-stimulatory receptors. However, more investigation is required to unravel and better understand how hypoxia influences immune cells. For instance, HICs could be used to study DC-T cell interactions in a hypoxic environment or understand how the ECM components synergize with hypoxia to inhibit anti-cancer immune responses.(*76*) In addition, our study displayed several limitations, such as the use of only one end point, one O_2_ tension in our observations, and an artificial B16-OVA/OT-1 T immunology model. Investigating the short and long-term impact of hypoxia and developing more accurate HIC-based cancer immunology models would have the potential to deepen our understanding of the role of hypoxia on immunosuppression and immune cell function across various key immune cells such as NK cells and macrophages.

In summary, we engineered the first injectable hypoxia-inducing and macroporous biomaterial capable of emulating solid tumor hypoxia for cancer and cancer immunology modeling. HICs can quickly induce hypoxic conditions and maintain hypoxia for days while remaining cytocompatible and biocompatible. Additionally, we report that HIC-supported cancer models recapitulate hypoxia-mediated metabolic switch and phenotypic changes toward aggressiveness, which was validated with resistance to clinically relevant cancer therapeutics. Finally, HICs can reproduce the immunosuppression of DCs and cytotoxic T cells when subjected to hypoxic TMEs. This allowed us to identify the implication of HA in retaining the cDC subset while altering the capacity of DCs to be activated and undergo maturation under hypoxic conditions. Importantly, these findings bring to light a new TME-mediated immunosuppression mechanism that was not previously reported. In addition, the HIC platform confirmed the key role of CTLA-4 over PD-L1-driven immunosuppression in solid tumor, suggesting that targeting NK cells to prevent cytotoxic T-cell elimination may be a promising avenue to strengthen tumor rejection. Considering that 97% of oncology clinical trials fail to receive FDA approval mostly due to poor drug efficacy,(*102*) the HIC platform has the potential to improve the efficiency of cancer drug screening and discovery, deepen our understanding of cancer immunology and unravel new mechanisms of anti-tumor immune responses, and ultimately advance immunotherapy for clinical and translational research.

## Methods

### Materials

HA sodium salt PEG (10kDa), DPBS, sodium bicarbonate (NaHCO_3_), antibiotic/antimycotic solution, glycidyl methacrylate (GM), GOX (100–250 kDa U/g from Aspergillus niger, Type X-S), tetramethylethylenediamine (TEMED), ammonium persulfate (APS), Pierce^TM^ quantitative peroxide assay, Glucose enzymatic colorimetric assay (GAGO20), fetal bovine serum (FBS), paraformaldehyde (PFA), Triton X-100, TPZ, and 4′,6-diamidino-2-phenylindole (DAPI) were purchased from MilliporeSigma (St. Louis, MO). 5/6-carboxy-tetramethyl-rhodamine succinimidyl ester (NHS-rhodamine), 5/6-carboxyfluorescein succinimidyl ester (NHS-fluorescein), cyanine 5 succinimidyl ester (NHS-Cy5), penicillin, streptomycin, gentamycin, rabbit anti-mouse IgG (H+L) Alexa Fluor 488 secondary antibody, PureLinkTM RNA Mini Kit, High Capacity cDNA Reverse Transcription Kit gene expression assays, pacific green fixable Live/Dead, and alamarBlue™ were purchased from Thermo Fisher Scientific (Waltham, MA). CAT (2,000 U/mg, Aspergillus niger) was purchased from MP Biomedical (Irvine, CA). APC-conjugated anti-mouse HIF-1α (Clone 241812), IFN-γ, TNF-α, IL-6, and granzyme B DuoSet ELISA kits were purchased from R&D Systems (Minneapolis, MN). OVA 257-264 (SIINFEKL peptides), CpG ODN 1826 (ODN, 5’-tccatgacgttcctgacgtt-3’, VacciGrade) and LPS from E. coli 0111:B4 (LPS-EB VacciGrade) were purchased from InvivoGen (San Diego, CA). Amine-terminated GGGGRGDSP (G_4_RGDSP) peptide was purchased from Peptide 2.0 (Chantilly, VA). Acrylate-PEG-N-hydroxysuccinimide (AP-NHS, 3.5 kDa) was purchased from JenKem Technology (Plano, TX). ViaQuant™ Fixable Far-Red Dead Cell Staining Kit was purchased from GeneCopoeia (Rockville, MD). Alexa Fluor 488-phalloidin was purchased from Cell Signaling Technology (Danvers, MA). The Hypoxyprobe Plus kit was purchased from Hypoxyprobe Inc. (Burlington, MA). Lactate-Glo^TM^ and Glutamine/Glutamate-Glo^TM^ assay kits were purchased from Promega (Madison, WI). The perforin ELISA kit and Fixable Viability Dye eFluor 780 were purchased from eBioscience (St. Louis, MO). Dulbecco’s Modified Eagle Medium (DMEM), Roswell Park Memorial Institute 1640 medium (RPMI), 2-mercaptoethanol and L-glutamine were purchased from Gibco (Waltham, MA). Murine melanoma cells (B16-F10, CRL-6475) and breast cancer cells (4T1, CRL-2539) were purchased from ATCC (Rockville, MD). Microsep^TM^ Advance Centrifugal Devices with Omega Membrane 3K was purchased from Pall Corporation, (Port Washington, NY). DOX and CIS were purchased from Tocris Bioscience (Bristol, UK). mGM-CSF was purchased from GenScript (Piscataway, NJ). The following reagents were all purchased from Biolegend (San Diego, CA): PE-conjugated anti-mouse CD317 (clone 927), PerCP/Cyanine5.5-conjugated anti-mouse major histocompatibility complex (MHC)-II (I-A/I-E, clone M5/114.15.2), PerCP/Cyanine7 anti-mouse F4/80 (clone BM8), APC-conjugated anti-mouse CD11c (clone N418), Alexafluor 700-conjugated anti-mouse CD86 (Clone GL1), APC/Cyanine7 anti-mouse/human CD45R/B220 (clone RA3-6B2), Alexa Fluor 488-conjugated anti-mouse MHC-I (Clone AF6-88.5), PerCP/Cyanine7-conjugated anti-mouse CD39 (Clone Duha59), APC/Cyanine7-conjugated anti-mouse CD73 (Clone TY/11.8), PE-conjugated anti-mouse CD137 (Clone 17B5), PerCP/Cyanine5.5-conjugated anti-mouse CD25 (Clone PC61), PerCP/Cyanine7-conjugated anti-mouse H-2Kb bound to SIINFEKL (Clone 25-D1.16), APC-conjugated anti-mouse CTLA-4 (Clone UC10-4B9), Alexa Fluor 700-conjugated anti-mouse CD8a (Clone 53-6.7), APC/Cyanine7-conjugated anti-mouse CD69 (Clone H1.2F3), BV401-conjugated anti-mouse PD-1 (Clone 29F.1A12), FITC-conjugated anti-mouse CD3 (Clone 145-2C11), BV605-conjugated anti-mouse CD44 (Clone IM7), pacific green fixable Live/Dead, BV650-conjugated anti-mouse LAG3 (Clone C9B7W), FITC-conjugated anti-mouse granzyme B (Clone: QA16A02), APC-conjugated anti-mouse perforin (Clone: S16009B), BV421-conjugated anti-mouse TNF-α (Clone: MP6-XT22), BV421-conjugated anti-mouse IFN-γ (Clone XMG1.2) brefeldin A, monensin, Cyto-Fast Fix/Perm Buffer Set.

### Cryogel and HIC fabrication

#### Polymer modification

HAGM (degree of methacrylation: 32%) and APR were prepared as previously described.(*39*) For the synthesis of APG and APC, AP-NHS was first reacted in NaHCO_3_ buffer solution (0.1 M, pH 8.5) for 4 h at RT with either GOX or CAT (molar ratio = 1:3), respectively. Next, the products (APG or APC) were freeze-dried overnight, and stored at −20 °C until further use.

#### Cryogelation

Cryogels were fabricated at subzero temperature as previously described(*32, 103*) Briefly, HAGM (4% w/v) and APR (0.8% w/v) were first dissolved in deionized water (dH_2_O). Next, the solution was precooled at 4°C, mixed with TEMED (0.07% w/v) and APS (0.28% w/v), poured into a Teflon® mold on ice (cubiform: 4 x 4 x 1 mm^3^, 16 μL/cryogel), and then transferred into a freezer at −20°C for 16 h to allow cryopolymerization. Finally, the newly formed cryogels were thawed at RT and washed several times with dH_2_O. For the fabrication of PEGDM cryogels, PEGDM was synthesized as previously reported.(*104*) Briefly, PEGDM (10% w/v) and APR (0.8% w/v) were dissolved in dH_2_O. Next, PEGDM cryogels were prepared similarly to the method described above. For HIC fabrication, APC and APG enzymes were used during cryogelation. Briefly, APC and APG were mixed with HAGM (4% w/v) or PEGDM (10% w/v) at various concentrations (0.1–1% w/v). Next, the polymer solution (HAGM/APC or PEGDM/APC) was precooled at 4°C, mixed with TEMED (0.07% w/v) and APS (0.28% w/v), poured into a Teflon® mold on ice (cubiform: 4 x 4 x 1 mm^3^, 16 μL/cryogel), and then transferred into a freezer at −20°C for 16 h to allow cryopolymerization. For confocal imaging, 0.3% of HAGM was substituted with rhodamine-labeled HAGM(*105*) in each polymer solution prior to cryogelation to fluorescently label the polymer network of cryogels.

### Physical characterization of HICs

The microstructural features of HICs were characterized by confocal microscopy using an LSM 800 (Zeiss, Oberkochen, Germany) or scanning electron microscopy using a Hitachi S-4800 scanning electron microscope (Tokyo, Japan) as previously described.(*39*) For assessing the enzyme grafting efficiency, APG and APC were labeled with NHS-fluorescein and NHS-Cy5, respectively, at a molar ratio of 1:15 (enzyme:fluorophore). For scanning electron microscopy, the cryogels were lyophilized for 24 h, mounted on a sample holder using carbon tape, sputter-coated with a 5 nm layer of platinum/palladium, and then imaged at a voltage of 25 kV and a current of 10 µA. Pore sizes were determined using Fiji,(*106*) Analyze Particles, and Diameter J software plugins.

The degree of pore connectivity was evaluated by a water-wicking technique assay. Hydrated HIC disks (6-mm diameter, 1-mm height) were first weighed on an analytical scale, then free water within the interconnected pores was wicked away using Kimwipe®, and the cryogels were weighed a second time. The degree of pore connectivity was calculated as the volume of water (%) wicked away from the scaffolds.

The swelling ratio was determined using a conventional gravimetric procedure.(*29*) Cylindrical HICs (6-mm diameter, 6-mm height) were fabricated, equilibrated in DPBS for 24 h at 37 °C, and weighed (m_s_). Next, the cylinders were washed in dH_2_O, lyophilized, and weighed again (m_d_). The equilibrium mass swelling ratio (Qm) was calculated as the mass of fully swollen HICs (m_s_) divided by the mass of lyophilized HICs (m_d_) or Qm= m_s_/m_d._

The mechanical properties of cryogels were determined by measuring their stiffness using an Instron 5944 universal testing system (Instron, Norwood, MA). Cylindrical HICs were dynamically deformed at a constant rate (strain rate of 10% per min) between two parallel plates for 10 cycles under continuous hydration in DPBS. Numerical values of the compressive strain (mm) and load (N) were then measured at the 8^th^ cycle on an Instron’s Bluehill 3 software. Young’s moduli (or Elastic moduli) were reported as the tangent of the slope of the linear (elastic) region on the stress-strain curve.

The syringe injectability of HICs was recorded or photographed using a Canon camera. Cuboidal cryogels (4 x 4 x 1 mm) were suspended in DPBS (0.2 mL), loaded into a 1 mL syringe, and subsequently injected through a 16-gauge hypodermic needle into a petri dish.

### Oxygen depletion kinetics

Oxygen tension in HIC containing well plates were determined using optical O_2_ sensor spots (OXSP5, PyroScience GmbH, Aachen, Germany). OXSP5 were glued onto the bottom of 48-well plate wells using silicone glue (SPGLUE, PyroScience) and dried for 16 h. DPBS (0.2 mL) supplemented with 10 g/L D-glucose and 1% antibiotic/antimycotic solution) was added to each well containing sensor spots and placed in a humidified incubator (37°C, 5% CO_2_) (Thermo Fisher Scientific). Once equilibrated, cuboidal HICs were individually placed into each well, and then the dissolved O_2_ concentration (mmHg or %, 1 point every 300 s) was recorded for up to 6 days. DPBS was added to wells surrounding HIC-containing wells to prevent dehydration. A 10X solution (100 g/L D-glucose, 10% antibiotic/antimycotic solution) was added into each cryogel-containing well every 2 days (20 µL). In these studies, normoxia was set as 140 mmHg or 18.6% O_2_, whereas hypoxia was set as 7.4 mmHg or 1% O_2_.

### Hydrogen peroxide production and glucose consumption

The production of H_2_O_2_ by HICs in cell culture was examined using a Pierce^TM^ quantitative peroxide assay. Cuboidal HICs formulated at various APC concentrations (0.1–1%) were incubated into 24-well plates containing 0.6 mL of DPBS supplemented with 4.5 g/L D-glucose and 1% antibiotic/antimycotic solution for 6 days at 37°C. Every day, 100 µL of the supernatant was collected from each well and replaced with fresh DPBS. The supernatants were spin filtered using Microsep^TM^ Advance Centrifugal Devices with Omega Membrane 3K (4000 G, 4°C, 45 min), and subsequently characterized with the peroxide assay.

The kinetics of glucose consumption were determined using a glucose enzymatic colorimetric assay. Cuboidal HICs formulated at various APG concentrations (0.01–0.1%) were incubated into 48-well plates (4 cryogels/well) containing 0.8 mL of DPBS supplemented with 6 g/L D-glucose and 1% antibiotic/antimycotic solution for 6 days at 37°C. A total of 25 µL of the supernatants from each well was collected daily and quantified for its glucose content using a glucose assay kit (GAGO20).

### Cell seeding and culture

B16-F10 and 4T1 cells were cultured in complete DMEM (DMEM supplemented with 10% FBS, 100 µg/mL penicillin, and 100 µg/mL streptomycin). B16-F10 ovalbumin (OVA)-expressing cells (B16-OVA, kindly provided by Prof. Sitkovsky at Northeastern University) were cultured in complete DMEM supplemented with 50 µg/mL gentamycin. Cells were incubated at 37°C in either humidified 5% CO_2_/95% air (normoxia) or humidified 5% CO_2_/1% O_2_/94% N_2_ (hypoxia). Prior to cell seeding, cuboidal HICs were sanitized with 70% ethanol for 10 min, washed several times with DPBS, partially dehydrated on a gauze under sterile conditions, and then placed in 48-well plates. Finally, a cell suspension (B16-F10, B16-OVA, or 4T1) in complete DMEM (10^7^ cells/mL, 10 µL) was added dropwise onto each cryogel and incubated for 1 h to allow cell adhesion. The cell-laden HICs were then supplemented with additional complete DMEM (0.5 mL/well) for the duration of the experiment. For experiments requiring a hypoxic environment, complete DMEM was preconditioned for 24 h under hypoxic conditions.

### Cell viability

Cell viability of cell-laden HICs was evaluated by a live/dead assay. Following a 24 h incubation period, cell-laden HICs were incubated for 15 min with a ViaQuant™ Fixable Far-Red Dead Cell Staining Kit according to the manufacturer’s instructions. Next, HICs were washed twice with DPBS, fixed with 4% PFA for 20 min at RT, and then rinsed two more times with DPBS. Next, cells were permeabilized with 0.1% Triton X-100 in DPBS for 15 min, stained with Alexa Fluor 488-phalloidin and DAPI following the manufacturer’s protocols, and finally imaged by confocal microscopy (LSM 800). Tile images were recreated from five cell-laden HICs per given condition to image their dimensions in their entirety. Cell viability, quantified using the Fiji software, was defined as the ratio between the total number of dead cells (i.e., cells stained in red) and the total number of cells (i.e., cell nuclei stained in blue and green fluorescence). For each condition, one representative stacked image (4096 x 4096 pixels) was captured, and slices were separated by 6μm throughout the z-stack (20 stacks).

### Hypoxia detection

Cellular hypoxia was determined using a Hypoxyprobe^TM^-1 kit according to the manufacturer’s recommendations. After 24 h of incubation in normoxic conditions, 50 μL of 2 mM Hypoxyprobe^TM^-1 (200 µM final) was added to cell culture media and incubated for 2h. Next, the cells were fixed (4% PFA), permeabilized (0.1% Triton X-100 in DPBS), and then immunostained with mouse Mab1 antibody, rabbit anti-mouse IgG (H+L) Alexa Fluor 488 secondary antibody, and DAPI. Finally, the cell-laden cryogels were imaged by confocal microscopy (LSM 800). Fractions of cellular hypoxia, quantified using the Fiji software, was defined as the ratio between the total number of hypoxic cells (i.e., cells labeled in magenta) and the total number of cells (i.e., cell nuclei labeled in blue).

The gene expression of HIF-1α, vascular endothelial growth factor α (VEGFα), cluster of differentiation 44 (CD44), CD73, SRY-Box Transcription Factor 2 (SOX2), and hypoxanthine guanine phosphoribosyl transferase (HPRT) were assessed by quantitative reverse transcription polymerase chain reaction (RT-qPCR, Mx3005P QPCR System (Agilent, Santa Clara, CA). After 24 or 48 h of incubation in normoxia, total RNA was extracted from cell-laden cryogels using a PureLink^TM^ RNA Mini Kit per the manufacturer’s instructions. For RT-qPCR, mRNA retro-transcription was performed using a High-Capacity cDNA Reverse Transcription Kit on a MyCycler (Bio-Rad, Hampton, NH). Next, gene expression was quantified using the following TaqMan® Gene Expression Assays on an Mx3005P QPCR System: HIF-1α: Mm00468869_m1, VEGFα: Mm00437306_m1, CD73: Mm00501915_m1, SOX2: Mm03053810_s1, and HPRT (housekeeping gene): Mm03024075_m1.

### Cell metabolism

The anaerobic metabolism was evaluated by measuring the pH, but also the lactate, glutamine, and glutamate levels within the cell culture media of cell-laden HICs and blank cryogels (control) for 72 h. To determine the pH, 100 μL of the supernatant was collected every day and analyzed using a PH8500 portable pH meter (Southern Labware, Cumming, GA). To quantify lactate, glutamine, and glutamate concentrations, 100μL of the supernatant was collected daily and subsequently tested using Lactate-Glo^TM^ and Glutamine/Glutamate-Glo^TM^ assays per the manufacturer’s recommendations.

### Biocompatibility assessment

Seven-week-old female C57BL/6J mice (n = 5, The Jackson Laboratory, Bar Harbor, Maine) were used. Anesthesia was induced with isoflurane (4% for induction and 1% for maintenance) in 100% O_2_ using an inhalation anesthesia system (300 SERIES vaporizer, VSS, Rockmart, GA). Blank cryogels, HICs (0.4% APC, 0.04% APG), and APC-free HICs (0.04% APG) were suspended in DPBS (0.2 mL) and subcutaneously injected in both dorsal-flank regions of mice using a 16-gauge hypodermic needle. At days 7 and 28 mice were euthanized and the cryogels were explanted with the surrounding tissues. Next, each sample was fixed for 48 h in 4% PFA, embedded in paraffin, cryosectioned into 5-μm-thick slices, and then stained with hematoxylin and eosin (H&E) as well as Masson’s trichrome (MT) for standard histological evaluation (iHisto, Salem, MA).

### Cancer cell responses to chemotherapeutic agents

To assess acquired resistance to chemotherapy or increased sensitivity to a hypoxia-activated prodrug, the viability of cancer cells subjected to DOX, CIS, or TPZ treatment was assessed under normoxic conditions. B16-F10 and 4T1 cells were seeded in blank cryogels or HICs in 48-well plates and cultured in complete DMEM supplemented with various concentrations of DOX, CIS, or TPZ (0–31.6 μM). The media was changed daily, and cell viability was evaluated using alamarBlue^TM^ assay once a day for 3 days. The fractions of viable cells for each drug concentration were calculated and normalized to untreated cells. The half-maximal inhibitory concentration (IC_50_) was determined as the concentration of chemotherapeutic agents killing 50% of cancer cells using the tangent of the slope of the linear region on the viability-concentration curves. After 72 h of incubation, B16-F10 and 4T1-laden cryogels and HICs treated with 0 or 2μM of DOX were assessed for cell viability and imaged by confocal microscopy as previously described.

### BMDC generation and in vitro DC activation assays

DC activation studies were performed using BMDCs isolated from 6-8-week-old female C57BL/6 mice (The Jackson Laboratory) as previously described.(*32*) Femurs and tibialis of mice were explanted, sanitized in 70% ethanol for 5 min, washed in DPBS, and then the bone marrow was flushed with DPBS (2 mL, 27G needle). Next, cells were dissociated by pipetting, centrifuged (5 min, 300 g), and resuspended (10^6^ cells/mL) in RPMI supplemented with 10% heat-inactivated FBS, 100 U/mL penicillin, 100 μg/mL streptomycin, 2 mM L-glutamine, and 50 μM 2-mercaptoethanol. At day 0, BMDCs were seeded in non-treated 6-well plates (2 x 10^6^ cells/well) in 5 mL of complete RPMI supplemented with 20 ng/mL mGM-CSF. At day 3, an additional 5 mL of RPMI supplemented with 20 ng/mL mGM-CSF was added to each well. At days 6 and 8, half of the media (5 mL) was removed and replaced with fresh RPMI supplemented with 10 ng/mL mGM-CSF. At day 10, BMDCs were collected and used.

BMDC activation assays were performed in 48-well plates containing complete 0.4 mL of RPMI supplemented with 10 ng/mL mGM-CSF. This study was conducted at 37 °C under normoxic (humidified 5% CO_2_/95% air) or hypoxic (humidified 1% O_2_/5% CO_2_/94% N_2_) conditions. First, BMDCs (2 x 10^6^ cells/well) were pre-incubated for 24 h in normoxia or hypoxia, and in presence of one gel system/well (blank cryogels or HICs). To induce activation, the RPMI was supplemented with CpG (1 µg/mL) or LPS (100 ng/mL) and BMDCs were cultured for 24 h. The negative control consisted of BMDCs cultured in RPMI supplemented with 10 ng/mL mGM-CSF. The state of BMDC stimulation and maturation was first evaluated by flow cytometry (Attune NxT flow cytometer, Thermo Fisher Scientific) using the following fluorescent antibodies: PE-conjugated anti-mouse CD317 (clone 927), PerCP/Cyanine5.5-conjugated anti-mouse MHC-II (I-A/I-E, clone M5/114.15.2), PerCP/Cyanine7 anti-mouse F4/80 (clone BM8), APC-conjugated anti-mouse CD11c (clone N418), Alexafluor 700-conjugated anti-mouse CD86 (Clone GL1), APC/Cyanine7 anti-mouse/human CD45R/B220 (clone RA3-6B2), and pacific green fixable Live/Dead. Next, the concentration of secreted proinflammatory cytokines (IL-6 and TNF-α) in the supernatant was quantified by ELISA. When HA-coating was required, a solution of HA (1mg/mL) in DPBS was deposited on each well plate bottom (0.2 mL/well) of a 48-well plate, incubated overnight at 4°C, and subsequently washed 3 times with DPBS prior to starting the activation studies with BMDCs.

### In vitro cytotoxicity assay in a B16-OVA melanoma model

B16-OVA cells (1 x 10^5^ cells/cryogel) were first seeded into cuboidal blank cryogels or HICs, then cultured in 48-well plates supplemented with RPMI, and subsequently incubated for 24 h under normoxic (∼ 21% O_2_) or hypoxic (∼ 1% O_2_) conditions). Splenocytes (2 x 10^5^ cells/well) isolated from OT-1 RAG mice (stock #003831, 8-week old C57BL/6-Tg(TcraTcrb)1100Mjb/J, The Jackson Laboratory) were cultured for 24 h under normoxic conditions. Next, splenocytes were prestimulated with SIINFEKL peptide (2 μg/mL) for 24 h in RPMI supplemented with 50 µM 2-mercaptoethanol. Once the splenocytes were activated, B16-OVA cells and splenocytes were cultured together for 24 h at a target/effector ratio = 1:2. At the end of the coculture study: (*i*) aliquots of cell culture media were collected to quantify by ELISA the concentrations of IFN-γ, TNF-α, granzyme B, and perforin; (*ii*) the cytotoxicity of spleen-derived OVA-specific (OT-1) T cells against B16-OVA was assessed by a fixable Live-dead viability assay; and (*iii*) the expression of activation, maturation, and inhibition markers on B16-OVA and OT-1 T cells were evaluated by flow cytometry. The following fluorescent antibodies were used. For B16-OVA cells: Alexa Fluor 488-conjugated anti-mouse MHC-I (Clone AF6-88.5), PerCP/Cyanine7-conjugated anti-mouse CD39 (Clone Duha59), APC-conjugated anti-mouse HIF-1α (Clone 241812), APC/Cyanine7-conjugated anti-mouse CD73 (Clone TY/11.8), pacific green fixable Live/Dead, and BV650-conjugated anti-mouse PD-L1 (Clone 10F.9G2). For OT-1 T cells: Alexa Fluor 488-conjugated anti-mouse MHC-I (Clone AF6-88.5), PE-conjugated anti-mouse CD137 (Clone 17B5), PerCP/Cyanine5.5-conjugated anti-mouse CD25 (Clone PC61), PerCP/Cyanine7-conjugated anti-mouse H-2Kb bound to SIINFEKL (Clone 25-D1.16), APC-conjugated anti-mouse CTLA-4 (Clone UC10-4B9), Alexa Fluor 700-conjugated anti-mouse CD8a (Clone 53-6.7), APC/Cyanine7-conjugated anti-mouse CD69 (Clone H1.2F3), BV401-conjugated anti-mouse PD-1 (Clone 29F.1A12), pacific green fixable Live/Dead, and BV650-conjugated anti-mouse LAG3 (Clone C9B7W).

The cytotoxic activity of OT-1 T cells against B16-OVA cells was monitored by time-lapse microscopy. Briefly, B16-OVA cells (1 x 10^5^ cells/well) suspended in RPMI were seeded on the bottom of a 24-well plate for 3 h to allow adhesion. Next, 2 x 10^5^ activated splenocytes in RPMI (target/effector ratio = 1:2) were seeded on cuboidal blank cryogels or HICs and then transferred to each well (0.5 mL). Finally, time-lapse experiments were performed in a closed chamber using a bright field inverted microscope (Zeiss Axio Observer Z1, Zeiss, Oberkochen, Germany) and the coculture was conducted at 37 °C under normoxic conditions. One preselected region of each well (350 × 350 µm) was imaged every 10 min for 4 h.

To measure intracellular cytokine production, OT-1 T cells were cocultured with B16-OVA cells under normoxic conditions for 6 h within cuboidal blank cryogels or HICs in the presence of 1X Brefeldin A and 1X Monensin. Next, the cells were washed with DPBS, incubated at 4°C for 30 min with Fixable Viability Dye eFluor 780 in DPBS (1:1,000 dilution), washed twice with DPBS + 1% BSA (PBA), and to evaluate the expression of surface markers, cells were stained overnight at 4°C in PBA using the following antibodies: FITC-conjugated anti-mouse CD3 (Clone 145-2C11), Alexa Fluor 700-conjugated anti-mouse CD8a (Clone 53-6.7), and BV605-conjugated anti-mouse CD44 (Clone IM7). The next day, cells were washed thrice with PBA prior to being fixed and permeabilized using a Cyto-Fast Fix/Perm Buffer Set according to the manufacturer’s protocol. For the intracellular staining, cells were incubated in the permeabilization buffer for 30 min at 4°C with the following fluorochrome-conjugated antibodies: FITC-conjugated anti-mouse granzyme B (Clone: QA16A02), APC-conjugated anti-mouse perforin (Clone: S16009B), BV421-conjugated anti-mouse TNF-α (Clone: MP6-XT22), BV421-conjugated anti-mouse IFN-γ (Clone XMG1.2). Lastly, cells were washed thrice with the permeabilization buffer, resuspended in PBA, and then quantified by flow cytometry.

## Supporting information

Video 1 - Syringe injectability of 0.1% HICs.

Video 2 - Tumoricidal capacity of OT-1 T cells on B16-ova cells in normoxic conditions in the presence of blank cryogels

Video 2 - Tumoricidal capacity of OT-1 T cells on B16-ova cells in normoxic conditions in the presence of HICs.

## Acknowledgments

Authors thank the Institute for Chemical Imaging and Living Systems (ICILS) for their technical assistance with the confocal microscope. S.A.B. acknowledges support from the National Institutes of Health (NIH, 1R01EB027705 award) and National Science Foundation (NSF CAREER award, DMR 1847843).

## Author contributions

T.C., Z.J.R, and S.A.B. conceived and designed the experiments. T.C., Z.J.R., J.S., L.G., and M.H. performed the experiments. T.C., Z.J.R., and S.A.B. analyzed the data and wrote the manuscript. T.C., Z.J.R., and S.A.B. conceived the figures. All authors discussed the results, commented on and proofread the manuscript. The principal investigator is S.A.B.

## Supplementary Information

**Supplementary Figure 1:**
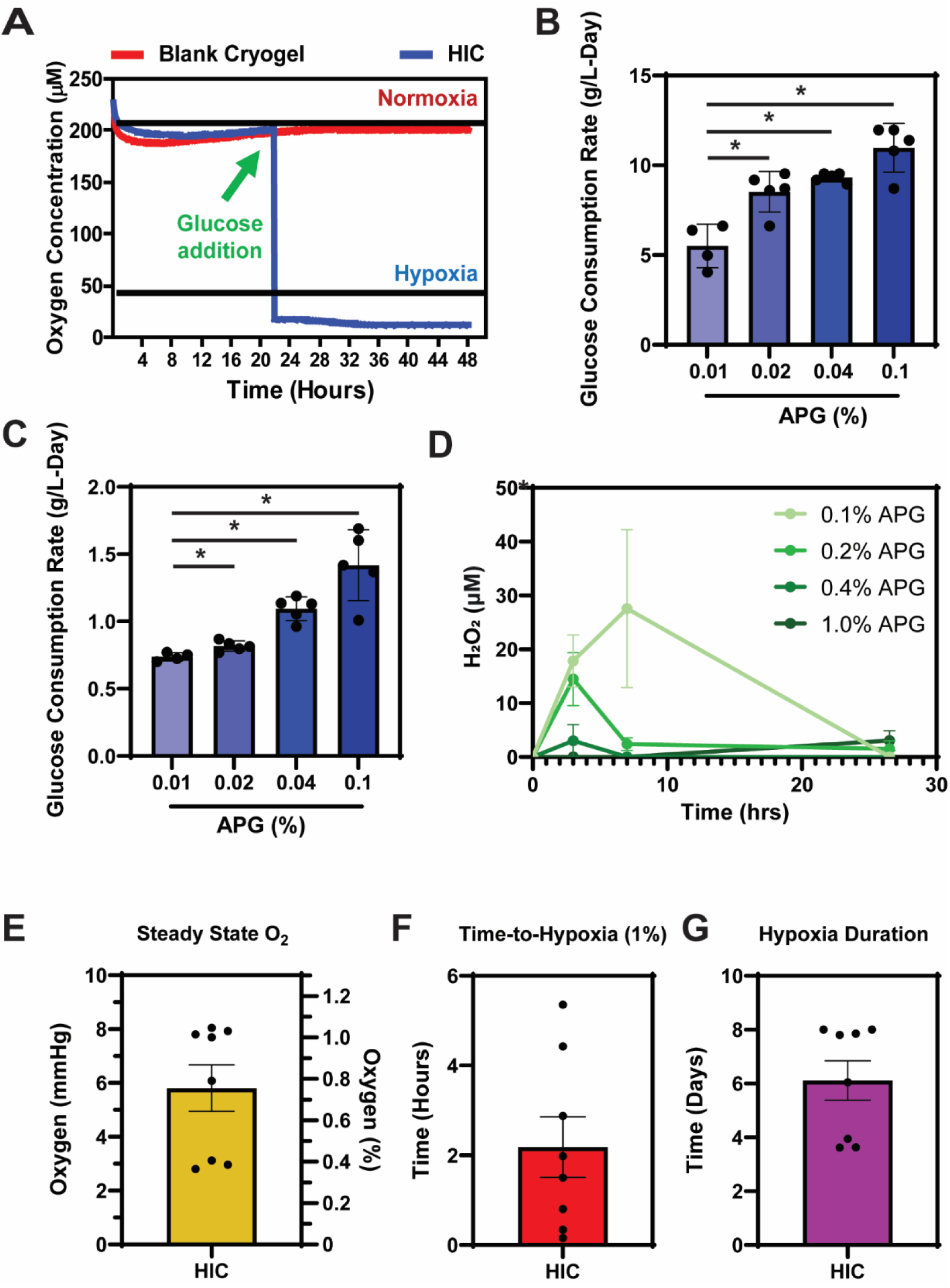
Glucose-dependent oxygen depletion induced by HICs. **A**) 0.04% HICs induce hypoxia following glucose addition to the medium. **B–C**) Glucose consumption rates of HICs containing various concentrations of APG (0.01–0.1%) during the induction of hypoxia (B) or the steady state phase (C). **D**) Hydrogen peroxide (H_2_O_2_) production from HICs during the induction of hypoxia. **E–G**) steady state of O_2_ (E), time-to-hypoxia (F), and hypoxia duration (G) induced by HICs (containing 0.4% APC + 0.04% APG). Values represent the mean ± SEM (n = 5 cryogels). Data were analyzed using ANOVA and Dunnett’s post-hoc test (* compared to 0.01% APG). **p < 0.05.

**Supplementary Figure 2:**
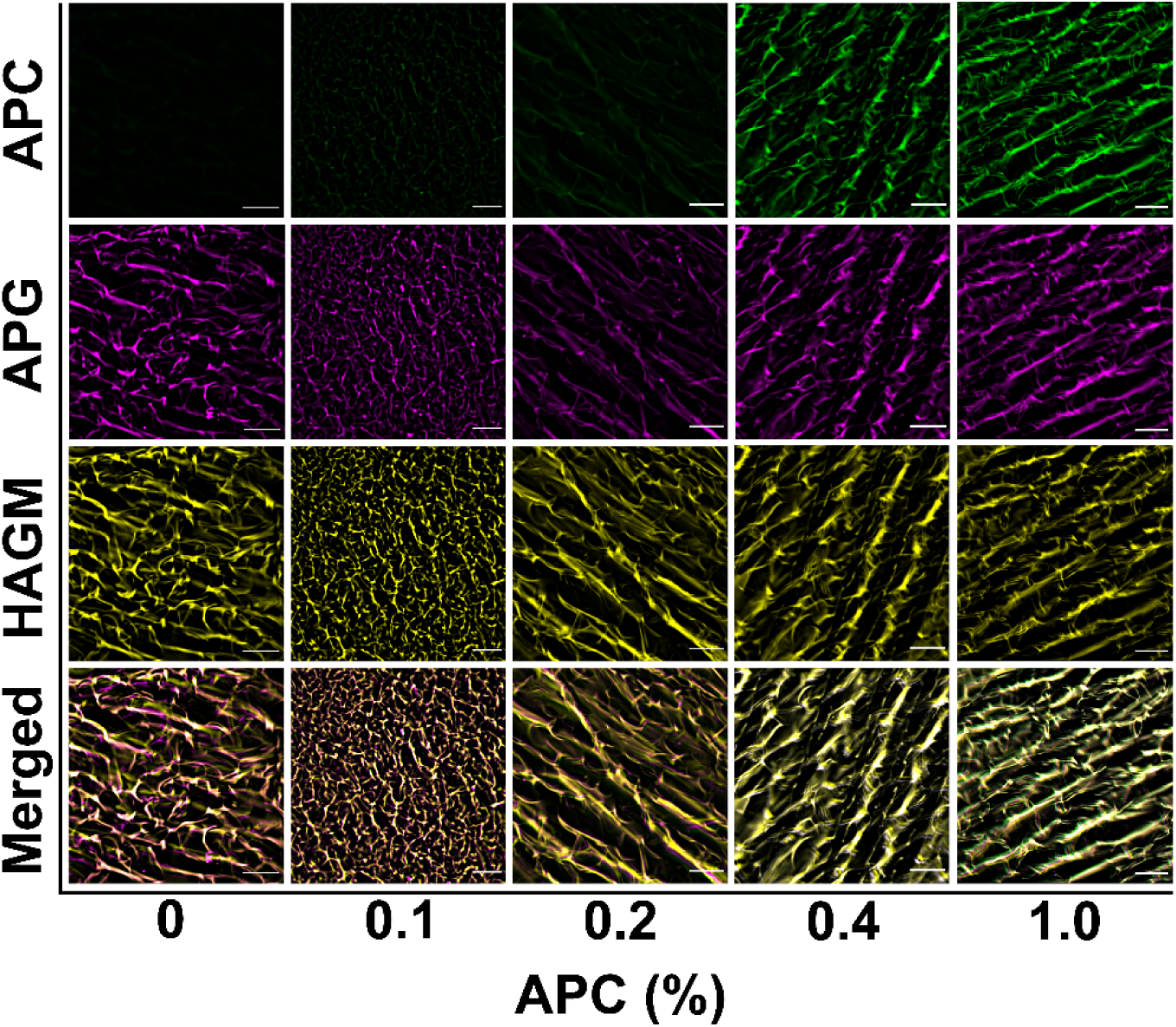
APC grafting efficiency within HICs. Confocal images showing the grafting efficiency of various APC concentrations (0–1%) within the polymer walls of cryogels. APC was fluorescently labeled with fluorescein (green) and APG with cyanine-5 (purple). Each image is representative of n = 5 samples. Scale bars = 100 µm.

**Supplementary Figure 3:**
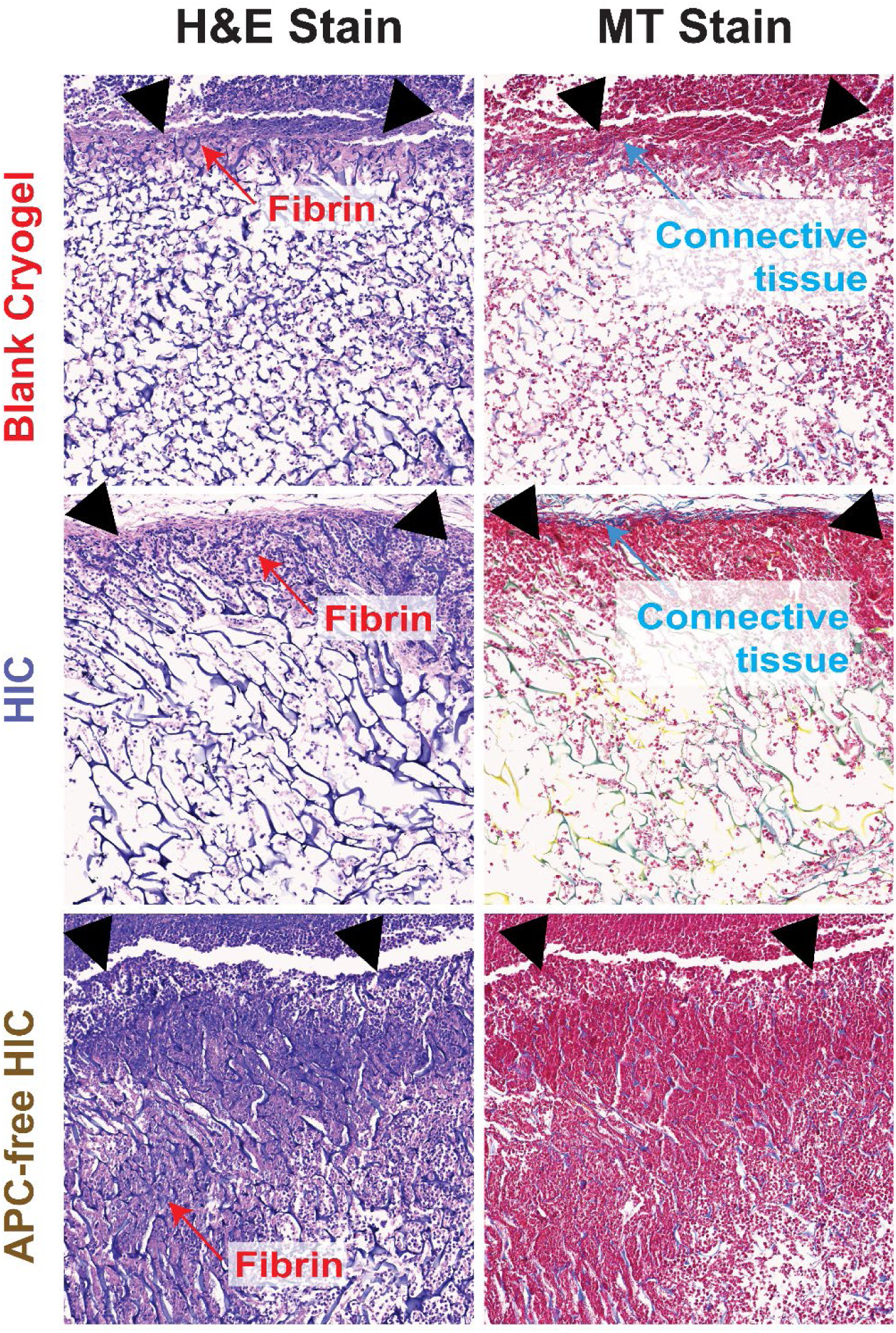
Cytocompatibility of HICs 3 days post subcutaneous injection. Histological analysis of explanted blank cryogels and HICs containing 0.04% APG and 0.4% APC (HIC) or only 0.04% APG (APC-free HIC). Hematoxylin and eosin (H&E) and Masson’s trichrome (MT) staining were performed. The arrows indicate the boundary between the cryogel and the host tissue. Histological images are representative of n = 5 samples per condition. Scale bars = 100µm.

**Supplementary Figure 4:**
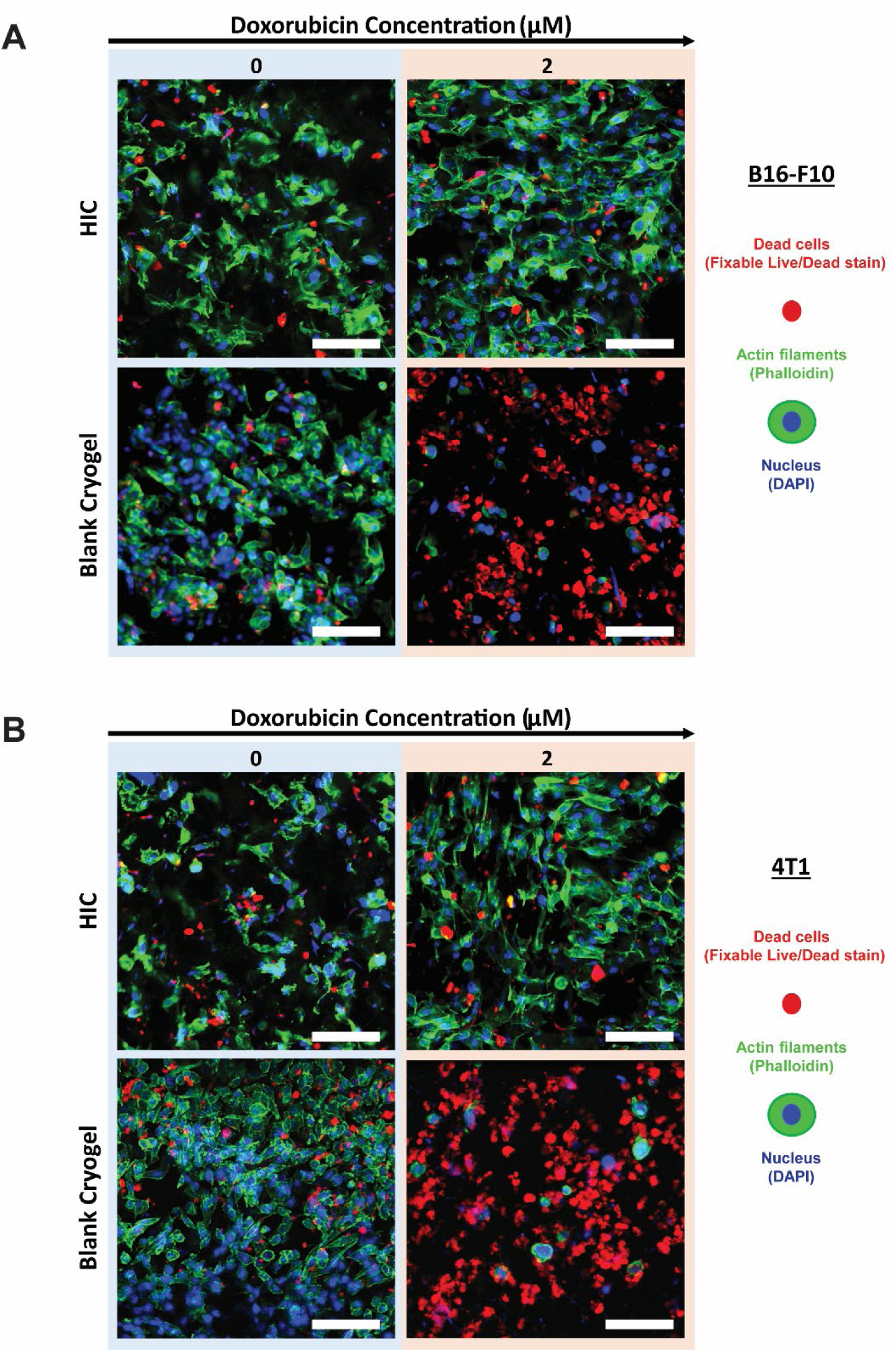
HICs induce resistance to chemotherapeutic treatments. **A–B**) Representative confocal images (n=5) of B16-F10 cells (A) and 4T1 cells (B) viability within blank cryogels and HICs after 72 h treatment with doxorubicin (2 μM) in normoxia. Blue = nuclei stained with 4′,6-diamidino-2-phenylindole (DAPI), red = dead cells stained with ViaQuant Far Red, green = actin cytoskeleton stained with Alexa Fluor 488 phalloidin, yellow = polymer walls stained with rhodamine. Confocal pictures are representative of n = 5 samples per condition. Scale bars = 100 μM.

**Supplementary Figure 5:**
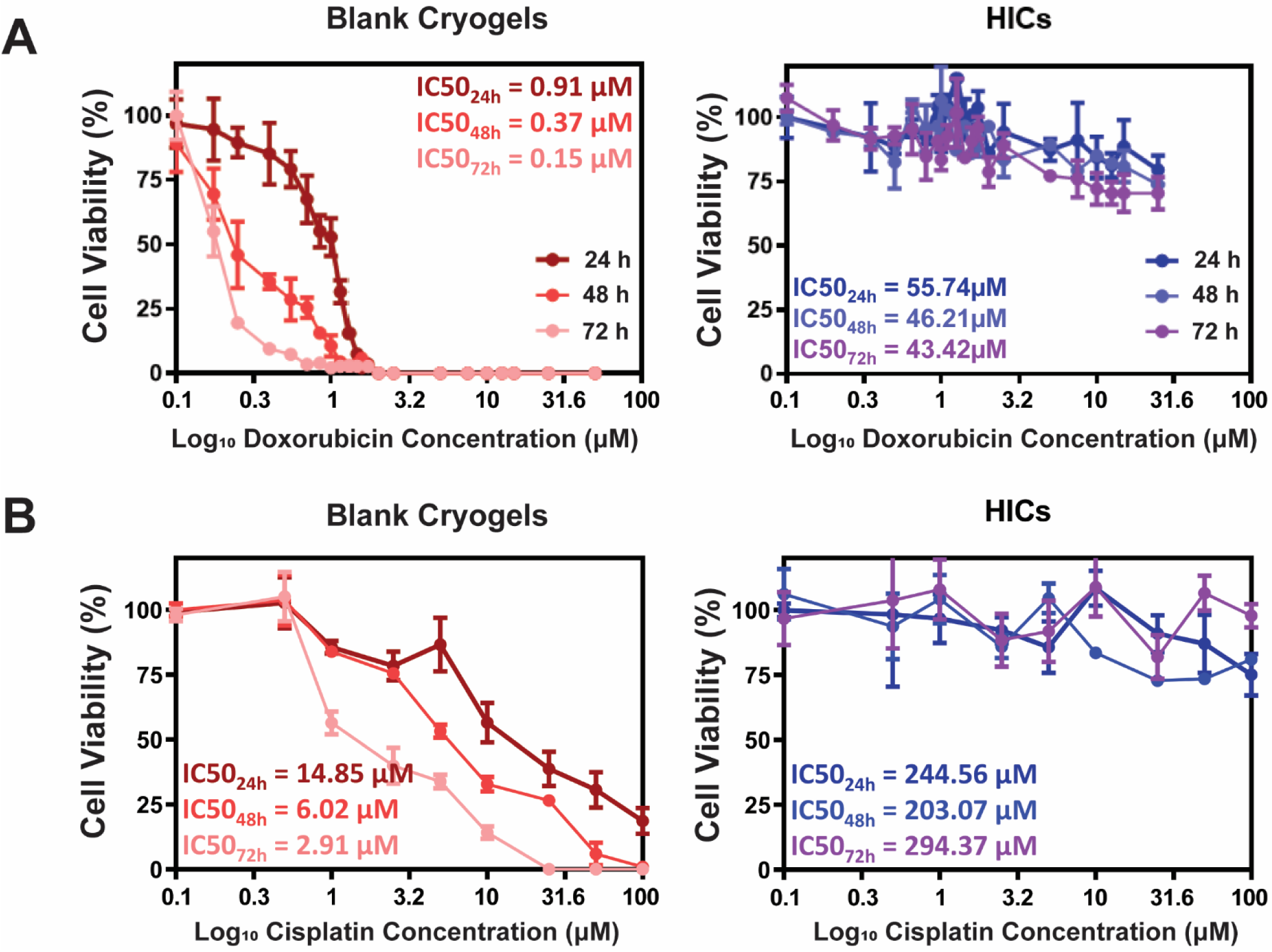
HICs inhibit doxorubicin and cisplatin-induced 4T1 cell death. **A–B**) 4T1 cell survival in blank cryogels and HICs after exposure to various concentrations (0–31.6 µM) of doxorubicin (A) and cisplatin (B) for 24, 48, and 72 h in normoxic conditions. Values represent the mean ± SEM (n = 5–6).

**Supplementary Figure 6:**
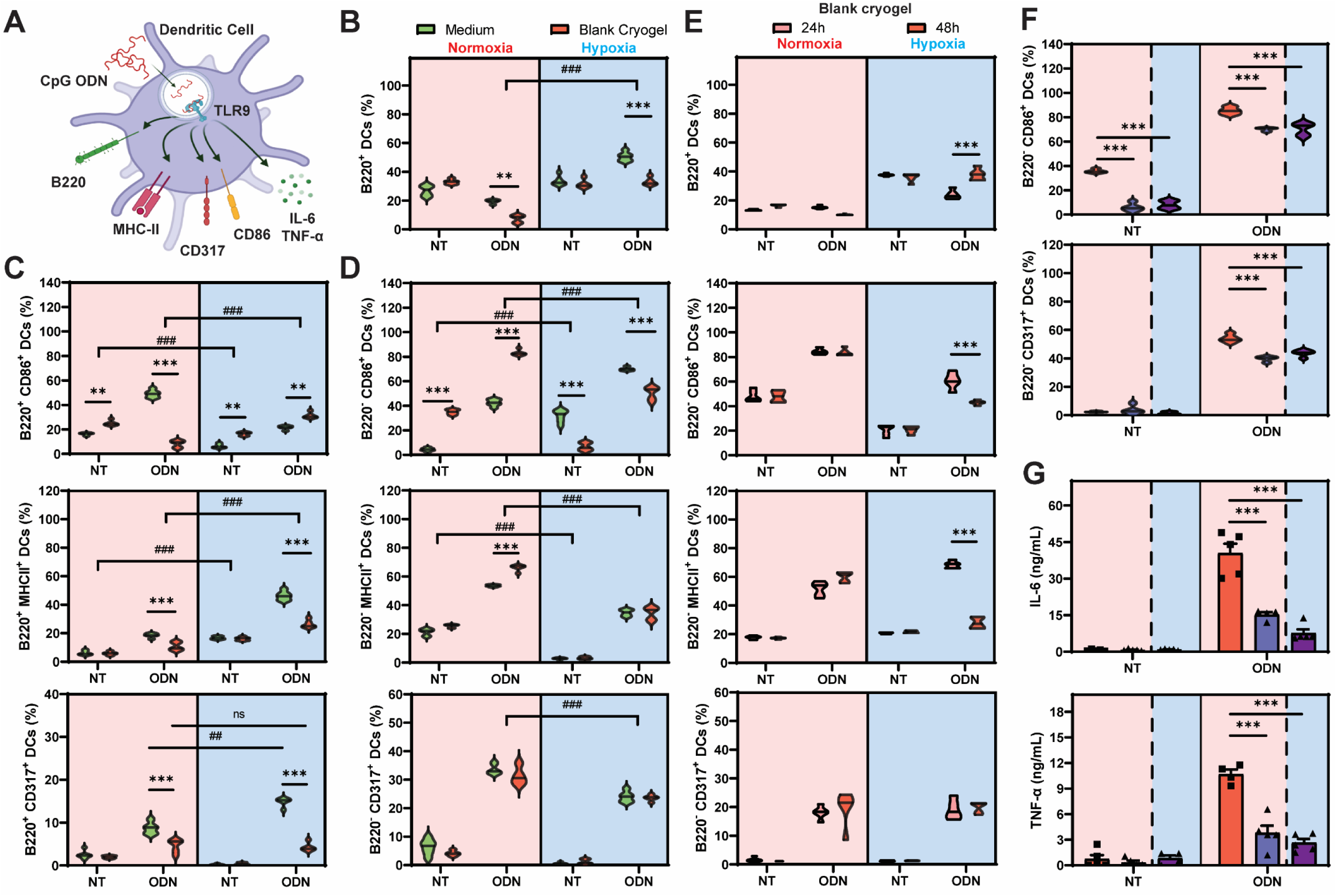
HICs impact DC subset and activation after CpG ODN stimulation. **A)** Schematic depicting DC activation in normoxia after CpG ODN stimulation. **B)** Fractions of B220^+^ CD11b^+^ CD11c^+^ after 48 h of incubation in normoxic or hypoxic conditions in medium alone or in the presence of blank cryogels. DCs were first pre-incubated for 24 h and then treated with LPS for 24 h. Non-treated DCs (NT) that received LPS-free medium were used as control. **C–D)** Fractions of activated B220^+^ (C) and B220^−^ (D) DCs determined using CD86, MHCII, or CD317 activations markers. **E)** Fractions of B220+ CD11b+ CD11c+ DCs, and of activated B220-DCs in the presence of blank cryogels without (24 h) or with (48 h) pre-incubation in normoxia or hypoxia. **F–G)** Evaluation of DC activation (F) and cytokine secretion (G) in the presence of blank cryogels or HICs. (F) Fractions of activated DCs were determined using CD86 or CD317 activations markers. (G) IL-6 and TNF-α concentration in the cell culture supernatant. Data were analyzed using ANOVA and Dunnett’s post-hoc test (*: compared to medium or blank cryogels, #: compared to normoxia). **p < 0.01, ^##^p < 0.01, ***p < 0.001, ^###^p < 0.001.

**Supplementary Figure 7:**
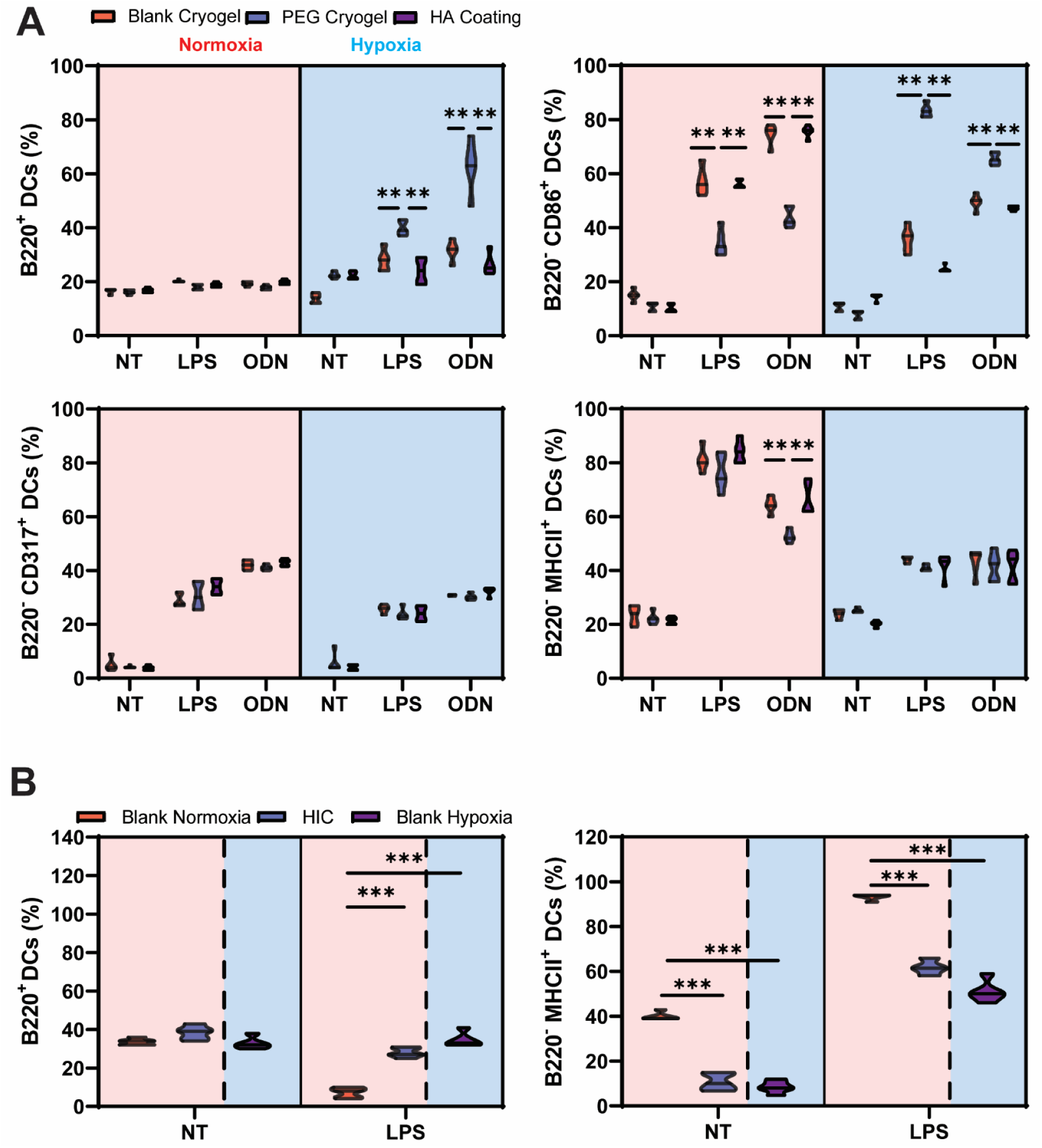
HICs recapitulate hypoxia-mediated DC inhibition in HA-rich environments. **A)** Fractions of B220^+^, B220^−^ CD86^+^, B220^−^ CD317^+^, and B220^−^ MHCII^+^ after 48 h of incubation in normoxic or hypoxic conditions well plates coated with HA or in the presence of blank (HAGM) cryogels or PEGDM cryogels. DCs were first pre-incubated for 24 h and then treated with LPS or ODN for 24 h. Non-treated DCs (NT) incubated in an LPS-free medium were used as a control. **B)** Fractions of B220^+^ DCs and B220^−^ MHCII^+^ DCs cultured in the presence of blank cryogels or HICs in normoxic or hypoxic conditions. Values represent the mean ± SEM (n = 5–6). Data were analyzed using ANOVA and Dunnett’s post-hoc test (compared to PEGDM (A) or blank (B) cryogels). **p < 0.01, ***p < 0.001.

**Supplementary Figure 8:**
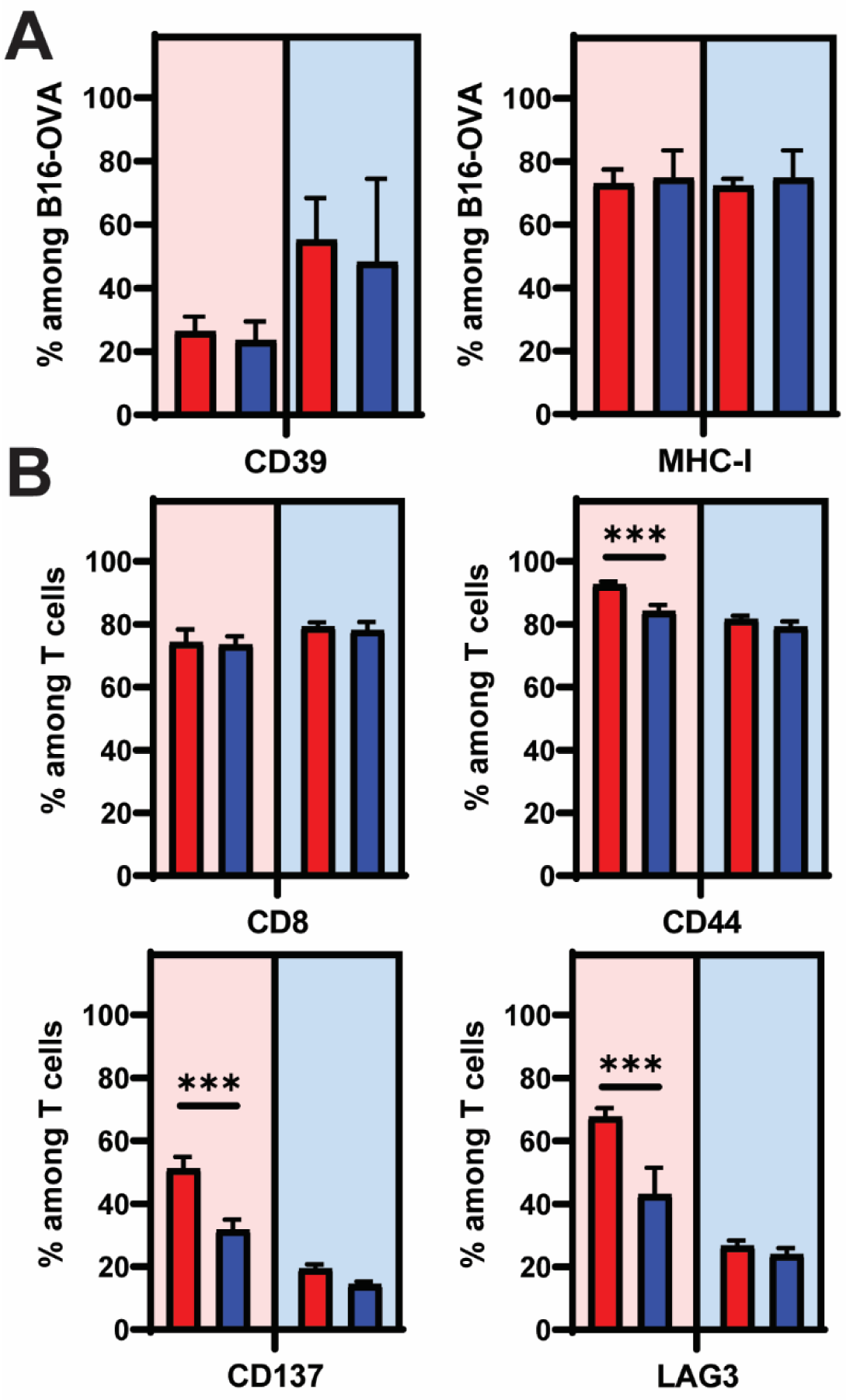
HICs impact T cell activation and function. B16-OVA-laden blank cryogels and OT-1 T cells were co-cultured in normoxic or hypoxic conditions for 24 h. **A)** Fractions of B16-OVA cells expressing CD39 and MHC-I. **B)** Fractions of OT-1 T cells expressing CD8, CD44, CD137, and LAG3. Values represent the mean ± SEM (n = 5–6). Data were analyzed using ANOVA and Dunnett’s post-hoc test (compared to blank cryogels). ***p < 0.001.

